# Sexually dimorphic role of estrogen receptor α in preserving right ventricular endothelial integrity

**DOI:** 10.1101/2025.11.20.689545

**Authors:** Jiajun Li, Vijaya Karoor, Andrea L. Frump, Hanqiu Zhao, Shunning Liang, Chen-Shan Chen Woodcock, Irina Petrache, Zhiyu Dai, Tim Lahm

## Abstract

Right ventricular (RV) function and adaptation to afterload increase determine survival in pulmonary hypertension (PH). RV adaptation in PH is sexually dimorphic and more preserved in females, mediated by protective estrogen receptor α (ERα) signaling in cardiomyocytes. However, the effects of ERα on RV endothelial cells (RVECs), a critical mediator of RV homeostasis and adaptation, are unknown. We hypothesized that ERα exerts sexually dimorphic pro-angiogenic effects on RVECs in vitro and promotes RV vascularization in vivo.

Compared to cells isolated from wild-type animals, RVECs from male and female rats with an ERα loss-of-function mutation (ERα^Mut^) showed reduced ability to form pseudo-vascular networks and migrate. RVECs from female ERα^Mut^ rats demonstrated increased apoptosis. In a PH model induced by monocrotaline (MCT), female ERα^Mut^ rats exhibited increased RV hypertrophy and reduced RV capillary density before (10 days) and at the time of established PH (28 days). Capillary rarefaction was associated with increased RVEC apoptosis, and, as identified by single-nucleus RNA-sequencing, by a net loss of the endocardial RVEC sub-population. Differentially expressed gene analysis and pathway analysis identified that capillary and endocardial RVECs from female MCT-PH ERα^Mut^ rats demonstrated decreased expression of migration pathways and increased expression of apoptosis pathways.

These findings reveal a sex-specific endothelial-intrinsic role of ERα that is essential for angiogenesis in the RV under both homeostatic and pathological conditions. This effect appears to stem from the enhanced survival and migration capacity of capillary and endocardial RVEC. Collectively, our results identify ERα as a potential target for developing sex-specific RV-directed therapies in PH.

**Translational perspective:** Effects of ERα on vascular function in RV failure induced by PH are poorly understood. We unveiled a novel sexually dimorphic role of ERα in regulating RV vascularization and RVEC function. Single nucleus RNA-Sequencing in female wild-type and ERα loss-of-function rats with PH identified 5 unique RVEC sub-populations under transcriptional control of ERα. Our findings provide insights into previously undescribed pro-angiogenic, pro-migratory and anti-apoptotic roles of ERα in female RVs and RVECs. Promoting RVEC migration or inhibiting RVEC apoptosis to enhance RV angiogenesis may be viable pathways to maintain RV function in PH patients of either sex. These findings offer novel opportunities and potential therapeutic avenues for preventing or treating RV failure.

## Introduction

Pulmonary hypertension (PH) is characterized by adverse vascular remodeling and intravascular pressure elevations in the lung^1^. However, right ventricle (RV) adaptation to the increased afterload is the major determining factor of survival^2–5^. Initially, the RV undergoes adaptive RV hypertrophy (RVH) to compensate for the pressure overload^6, 7^. However, as RV afterload continues to rise, maladaptive RVH develops, characterized by reduced RV ejection fraction as well as RV dilation and fibrosis^8, 9^. Pulmonary endothelial cell alterations have been one of the focuses in the pathogenesis of multiple types of PH, and pulmonary arterial endothelial cell dysfunction is a major contributor to lung vascular remodeling^10, 11^. However, the role of endothelial cell function in the RV has been less well studied.

While the pathophysiology of RV failure is complex and multi-factorial and involves a myriad of cell types, insufficient angiogenesis^12–15^ and impaired RV endothelial cell (RVEC) function^16^ have been identified as key contributors. Recent research revealed there is increased angiogenesis in the RV at early stages of PH-induced RVH^14,17^. This indicates that RVECs and RV capillaries may be able to adapt to RV pressure overload in early stages but fail to do so in later stages of RVH. However, mechanisms of RVEC adaptation and maladaptation to PH-induced RVH remain incompletely understood.

Importantly, RV adaptation in response to PH is sexual dimorphic, with female patients demonstrating better RV function^18^ and survival^19^. 17β-estradiol (E2), the most abundant female sex steroid, exerts RV-protective effects^16, 20–22^ that may include stimulation of angiogenesis^20, 22, 23^. Previous studies also suggest that estrogen receptor α (ERα) plays a major role in angiogenesis^23–26^. Importantly, ERα expression is reduced in RVECs from patients with PH-induced RV failure^16^. We recently demonstrated that ERα protects RV function in PH through increasing BMPR2 and apelin in RV cardiomyocytes^16^, both of which are known to be proangiogenic factors. However, it remains unknown if ERα regulates RVEC function and RV angiogenesis in the progression from adaptive RVH to RVF.

Here, we hypothesize that ERα exerts pro-angiogenic effects in the RV and in RVECs. We report that loss of ERα induces RVEC dysfunction and reduces angiogenesis both in vitro and in vivo. We identified an early onset of RVH and capillary rarefaction as well as increased RVEC apoptosis in female PH rats without functioning ERα. Single-nucleus RNA sequencing revealed newly identified RVEC subtypes in the female failing wild-type RV that exhibit impaired cell survival and migration signaling with the loss of ERα .

Our findings provide evidence for a sex-specific role of ERα in RV angiogenesis, with ERα specifically mediating protection against the development of RVH and RV capillary rarefaction.

## Methods

Main methods and techniques are presented here. Please see supplement for additional methods and information. All experiments were performed in accordance with recent recommendations^27, 28^, including randomization and blinding at the time of measurement and analysis.

### Monocrotaline (MCT)-PH

MCT (10 mg/ml in PBS; C2401, Sigma-Aldrich) was prepared on the day of injection. Male and female Sprague-Dawley rats or in-house bred wild-type (WT) and ERα loss-of-function mutant (ERα^Mut^) rats (200–250 g, 6-8 weeks of age) were subcutaneously injected with MCT (60 mg/kg) and maintained for 3, 10 or 28 days, followed by endpoint analysis.

### Hemodynamic assessment

Rats were anaesthetized by inhalation of 5% isoflurane an then orotracheally intubated and mechanically ventilated (100% Fi_O2_) during measurements with supplemental oxygen, after verification of euthermia, normocapnea, and normal pH. RV systolic pressure (RVSP) was measured by right heart cathetherization with 2-F Mikro-Tip® pressure catheter (Millar) via transjugular approach under 2% isoflurane as described previously^16^ and recorded with LabChart software (ADInstruments). Once all hemodynamic endpoints were obtained, animals were euthanized by exsanguination under anesthesia via the arterial line, followed by immediate organ harvest.

### RV hypertrophy assessment

Hearts were harvested and the atria were removed. The RV free wall was separated from the left ventricle (LV) and septum. The Fulton index was calculated as: RV weight / (LV + septum weight).

### Tube formation assay

RVECs were serum starved overnight (18h) with low-serum media (EBM phenored-free media with 0.5% charcoal stripped-FBS) prior to experimentation. Flat bottom 96-well plates (Corning) were coated with 40 µl of phenol red-free reduced growth factor Matrigel (Corning) per well, 1h before the experiment at 37°C in a humidified atmosphere containing 5% CO_2_. 10,000 RVECs in low-serum media were plated into each well for 24h. Plates were imaged at 10x magnification using Incucyte® Live-Cell Analysis System at 1h intervals. Images were then exported and uploaded to Wimasis.com for analysis.

### Transwell migration assay

RVECs isolated from WT and ERα^Mut^ rats were fserum-starved for 18h in low-serum media. Cells were then trypsinized and 50,000 RVECs were seeded to the top insert with 8µm pore size (Corning) in low-serum media (EBM phenored-free media with 0.5% charcoal stripped-FBS). Inserts were placed into a 24-well plate containing either low-serum medium (0.5% charcoal stripped-FBS) or low-serum medium supplemented with 10 ng/mL vascular endothelial growth factor A (VEGF-A; PeproTech) in the lower chamber to serve as a chemoattractant. Cells were incubated for 24 h at 37°C in a humidified incubator with 5% CO₂ to allow migration. Following incubation, inserts were washed twice with PBS and fixed in 4% paraformaldehyde for 10 min at room temperature. Non-migrated cells remaining on the upper surface of the membrane were gently removed with cotton-tipped applicators. Migrated cells on the lower surface were stained with 0.1% (w/v) crystal violet solution (Ward’s Science) for 20 min, rinsed with distilled water, and air-dried. Membranes were imaged under bright-field microscopy. Migrated cells were quantified in five random fields per insert using ImageJ software (NIH). Each condition was assayed in triplicate and repeated in at least three independent experiments.

### Wound healing assay

50,000 RVECs were seeded into each well of ImageLock 96-well plate (Sartorius). RVECs were then serum-starved overnight (18h) in low-serum media. A single layer of confluent RVECs was then scratched with Woundmaker Tool (Sartorius) and fresh low-serum media (EBM phenored-free media with 0.5% charcoal stripped-FBS) was added into each well with VEGF-A at a final concentration of 10 ng/ml. The ImageLock 96-well plate was then placed into Incucyte® S3 Live-Cell Analysis System (Sartorius) for imaging. 10x images of each well were taken every 2h. Duplicate samples from 3 different biological replicates were used. Migration was quantified as cells in initial wound area.

### Growth curve/cell counting assay

RVECs were seeded at a density of 2×10⁴ cells per well in 24-well plates containing 1 mL of EGM2-MV medium (Lonza) and allowed to adhere for 24 h at 37°C in a humidified atmosphere with 5% CO₂. After 24h, cells were serum-starved in low-serum medium (EBM phenored-free media with 0.5% charcoal stripped-FBS) for 48h. Following starvation, cells were detached using trypsin–EDTA (Gibco) and counted with a hemocytometer (Hausser Scientific); this time point was designated as day 0. At the same time, cell culture media was changed into either low-serum media or complete EGM2-MV media and cells were counted at days 1, 3, and 5 as described. Each condition was assayed in duplicate and repeated in at least three biological replicates.

### Caspase 3/7 activity assay

20,000 RVECs were plated into each well of the 96-well plate in 100µl EBM phenored free media (Lonza), and cells were allowed to attach for 24h at 37°C in a humidified atmosphere containing 5% CO_2_. 24h later, Caspase-Glo® 3/7 Assay (Promega) was performed on RVECs according to manufacturer’s protocol. Relative light units (RLU) were measured 30 min after the reagent was added.

### AnnexinV and PI assay

500,000 RVECs were plated into 10-cm dish in EGM2-MV media (Lonza). When confluency reached 70%, RVECs were serum-starved for 24h before proceeding to AnnexinV and PI assay (ProteinTech) using manufacturer recommened protocol. All stained RVECs were analyzed with LSRFortessa cell analyzer (BD Biosciences). Unstained cells were used as negative controls.

### Fluorescent and capillary staining

RV formalin-fixed, paraffin-embedded (FFPE) slides were rehydrated. Antigens were retrieved with citrate buffer in pressure cooker. Slides were cooled and incubated in PBS+Triton X-100 (0.25%) for 5min and washed with 3x PBST, 5 min each. Griffonia Simplicifolia Lectin (Vector Labs) was diluted in PBST at a final concentration of 5µg/ml. Slides were then incubated in Griffonia Simplicifolia Lectin at 4°C for 1 day. 1 day later, lectin was rinsed off with 3x PBST, 1 min each. Wheat Germ Agglutinin (WGA; Vector Labs) was diluted in PBST at a final concentration of 5 µg/ml. Slides were then incubated in WGA at room temperature for 2h. Lectin was then rinsed off with 3x PBST, 1 min each. 1 µg/ml of DAPI (Thermo Fisher Scientific) was used to stain the slides for 1 min to visualize nuclei and then rinsed off. Alides were dehydrated and mounted with ProLong™ Gold Antifade Mountant (Invitrogen). Slides were imaged with IX83 inverted microscope (Olympus).

### TUNEL (terminal deoxynucleotidyl transferase dUTP nick end labeling) assay

FFPE RV tissue sections were deparaffinized and rehydrated per the manufacturer’s protocol. Apoptotic cells were detected using In Situ Cell Death Detection Kit (Roche) following the manufacturer’s protocol. After completion of TUNEL labeling, slides were incubated overnight at 4°C with Griffonia simplicifolia lectin I (GSL I; Vector Laboratories) to visualize capillary structures. Slides were rinsed, mounted with ProLong™ Gold Antifade Mountant (Invitrogen), and imaged using an IX83 inverted fluorescence microscope (Olympus).

### Western blot analysis

Protein concentration was measured using BCA Protein Assay (ThermoFisher). Rat RV tissues (40 µg/sample) and RVECs (20 µg/sample) samples were prepared with 4x Laemmli Sample Buffer (Bio-Rad) and were run on 4–20% Mini-PROTEAN TGX Precast Protein Gels (Bio-Rad) in Tris/Glycine/SDS Running Buffer and transferred to Immobilon-P PVDF membranes (Millipore Sigma) using Tris/Glycine Running Buffer (Bio-Rad). The membranes were blocked with Pierce Protein-Free (TBS) Blocking Buffer (ThermoFisher) or 1% BSA in TBST. Primary antibodies (1:1000) were diluted in Blocking Buffer. Anti-rabbit-HRP and anti-mouse-HRP (Azure Biosystems) secondary antibodies were diluted 1:5000 in 1% BSA with TBST. SuperSignal™ West Femto Maximum Sensitivity Substrate (Thermo Scientific) was used to image the blots. Densitometry was performed using Image Lab (Bio-Rad). More detailed information on antibodies is listed in the supplement.

### Quantitative PCR

RVECs isolated from WT and ERα^Mut^ were first serum-starved for 18h in low-serum media. Total RNA was isolated from rat RVECs using RNeasy Plus Mini Kit (Qiagen). RNA concentration was measured by NanoDrop (Thermofisher). 1 µg total RNA was reverse-transcribed using iScript cDNA synthesis kit (Bio-Rad). Gene expressions were quantified with TaqMan assays. Changes in mRNA expression were determined by comparative CT (2^-ΔΔC^_T_) method.

### Single nucleus RNA-Seq (snRNA-Seq)

RV tissues from female WT and ERα^Mut^ rats were collected at 10 days and 4 weeks of MCT treatment. Nuclear extraction was performed on RV frozen tissues using 10x Genomics Chromium Nuclei isolation kit. cDNA libraries were prepared with GEM-X Universal 3’ Gene Expression v4 4-plex kit. Equal number of single cells isolated from WT and ERα^Mut^ rats was loaded on 10X Genomics Chromium Single Cell Controller to generate barcoded single cells for construction of single-cell cDNA libraries. SnRNA-seq libraries were sequenced on NovaSeq X Plus platform at Washington University in St. Louis Genomic Technology Access Center (GTAC). Cell Ranger (v9.0.0) was used for demultiplexing and counting. The rat reference genome Rattus_norvegicus_Rnor_6-0-101 was used as reference genome. R package Seurat (v5.3.0) was used for data preprocessing and visualization. Initially, cells with fewer than 100 or more than 4000 detected genes, or with >15% mitochondrial gene expression were removed. Genes expressed in <5 cells were discarded. Doublets were identified using a 2-layer approach as described previously: First, scDblFinder (v1.22.0) was used to predict potential doublets using default settings in an automated and unbiased fashion. Doublets were additionally identified manually when expressing combinations of marker genes from different cell types. After doublet removal, single cells were normalized with SCTransform and then used this method to integrate (https://satijalab.org/seurat/reference/integratedata).

For subclustering of ECs, clusters of interests were extracted. Resolution was set at 0.5. SingleR and Azimuth for human heart reference^29–31^ were then used for annotation.

Differential gene expression analyses between WT and ERα^Mut^ rat samples for each cluster of interest were performed using the Wilcoxon Rank-Sum test algorithm with log normalized counts in the RNA assay as input. A threshold of 0.25 for log fold change, 0.05 for the adjusted P value, and 0.1 for minimal fraction of cells was applied for downstream analysis.

Pathway analysis was performed with Qiagen Ingenuity Pathway Analysis software. Bar charts of top upregulated and downregulated pathways were selected within diseases and functions analysis. Pathways are ranked by activation Z-score.

### Data presentation

Due to the more pronounced phenotype observed in female rats, data from female rats are shown in the main manuscript. Data from male animals, due to space contraints, are shown in the supplement.

### Statistical analysis

Unless otherwise indicated, data represent mean ± SEM of biological replicates (run in technical triplicates). All statistical analyses were performed with Prism v9.0 (GraphPad Software, La Jolla, CA, USA). Sample sizes were estimated by power calculation. Correlations were determined using Pearson’s coefficient (R). For normally distributed data, comparisons between three or more groups were performed using two-way analysis of variance (ANOVA). For comparisons between two groups, significance was calculated with unpaired Student’s *t*-test. Statistically significant difference was accepted at p<0.05.

## Results

### ERα regulates RVEC angiogenic function and signaling in vitro

We first confirmed reduced ERα abundance in ERα^Mut^ rats. As shown previously^16^, female and male ERα^Mut^ rats exhibited an 80% reduction in RVEC expression of full-length *Esr1* transcript (**Supplementary Fig. 1A,B**). ERα^Mut^ rats also exhibited higher serum E2 levels than WT (**Supplementary Fig. 1C-F**).

We then sought to investigate whether ERα plays a pro-angiogenic role in RVECs. Compred to WT, loss of ERα resulted in reduced pseudo-vascular network forming capability (total tube length, numbers of branching points and rings) in both female (**Fig. 1A-D**) and male RVECs (**Supplementary Fig. 2A-D**). This suggests that ERα regulates pro-angiogenic function in RVECs. We next investigated effects of ERα on other significant contributors to angiogenesis, such as RVEC proliferation and migration^32–34^. Interestingly, female ERα^Mut^ RVECs exhibited increased proliferation compared to WT (**Fig. 1E**), whereas male ERα^Mut^ RVECs demonstrated reduced proliferation compared to their WT counterparts (**Supplementary Fig. 2E**). We next focused on RVEC migration. Both transwell migration and scratch assays revealed that loss of ERα resulted in decreased migration in female RVECs (**Fig. 2F-I**). Male ERα^Mut^ RVECs also exhibited decreased migration compared to WT RVECs (**Supplementary Fig. 2F-I**). These results suggest that ERα enhances pseudo-vascular network formation in RVECs in both sexes primarily via enhancing migration. ERα’s effects on RVEC proliferation, on the other hand, are sex-specific and appear to inhibit proliferation in female RVECs, indicating that impaired proliferation is unlikely to explain the reduced RVEC pseudo-vascular network formation found in ERα^Mut^ females.

**Figure 1.**
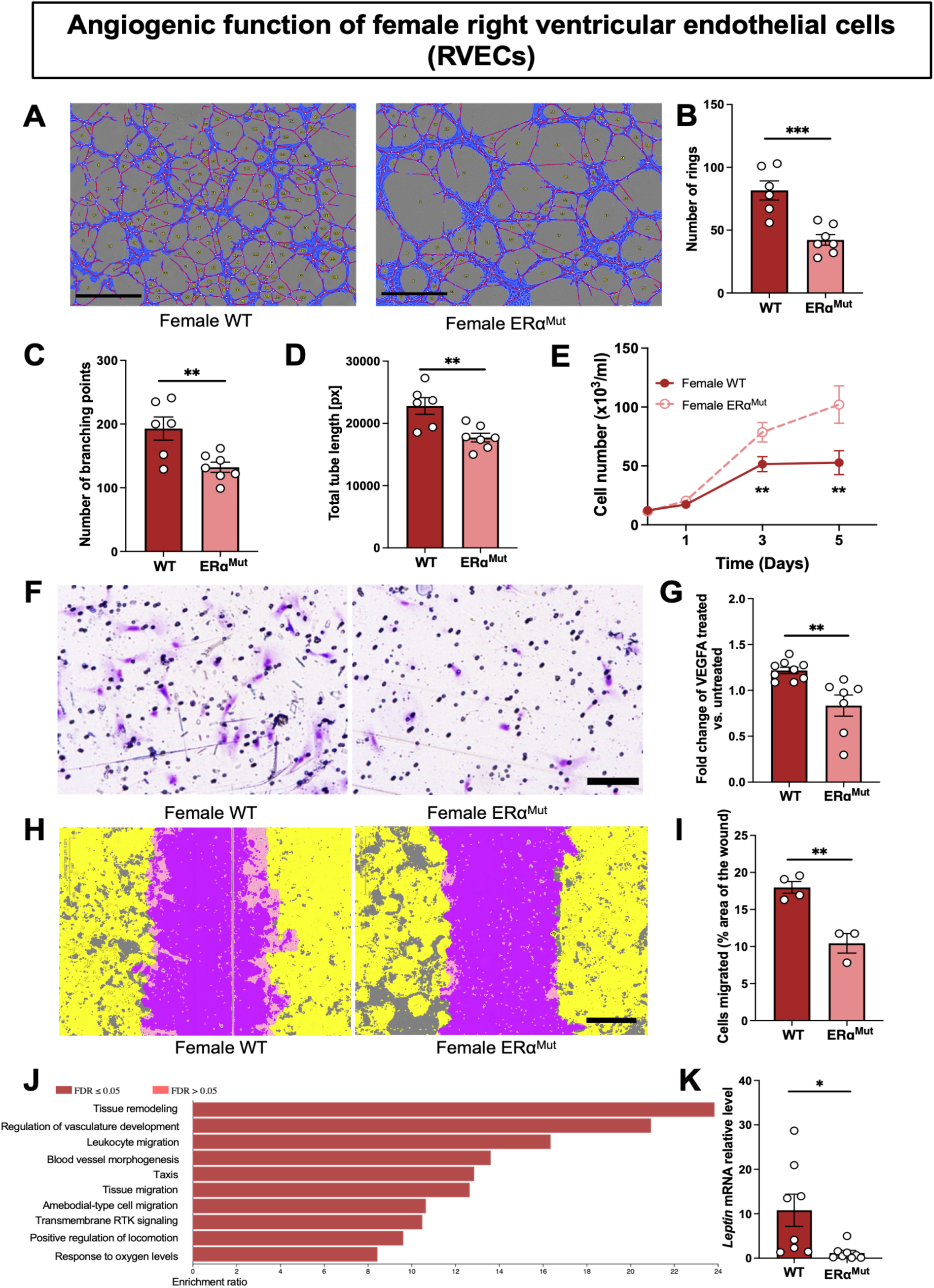
Loss of ERα reduces angiogenesis and migration in female right ventricle endothelial cells (RVECs). (**A-D**) Tube formation assays in female wild type (WT) and ERα^Mut^ RVECs at 10h after plating cells on Matrigel. Representative images (**A**) and quantification of number of rings (**B**), number of branching points (**C**) and total tube length (**D**). (**E**) Growth curves of female WT and ERα^Mut^ RVECs. (**F, G**) Transwell migration assay representative images and quantification of female WT and ERα^Mut^ RVECs at 24h. (**H, I**) Scratch assay representative images and quantification of female WT and ERα^Mut^ RVECs at 24h. Purple shade represents initial wound. (**J**) GO biological process comparisons between female WT and ERα^Mut^ RVECs from Qiagen rat angiogenesis microarray (n=4 animals/group, pooled). (**K**) qPCR analysis of leptin RNA expression of female WT and ERα^Mut^ RVECs from Taqman gene expression (relative expression level, 2-ΔΔCt calculation). Each data point = one animal. * p<0.05, ** p<0.01; *** p<0.001 by ANOVA with Tukey’s post-hoc analysis. Error bars represent mean ± SEM. Error bars represent mean ± SEM. Scale bars, 200µm

**Figure 2.**
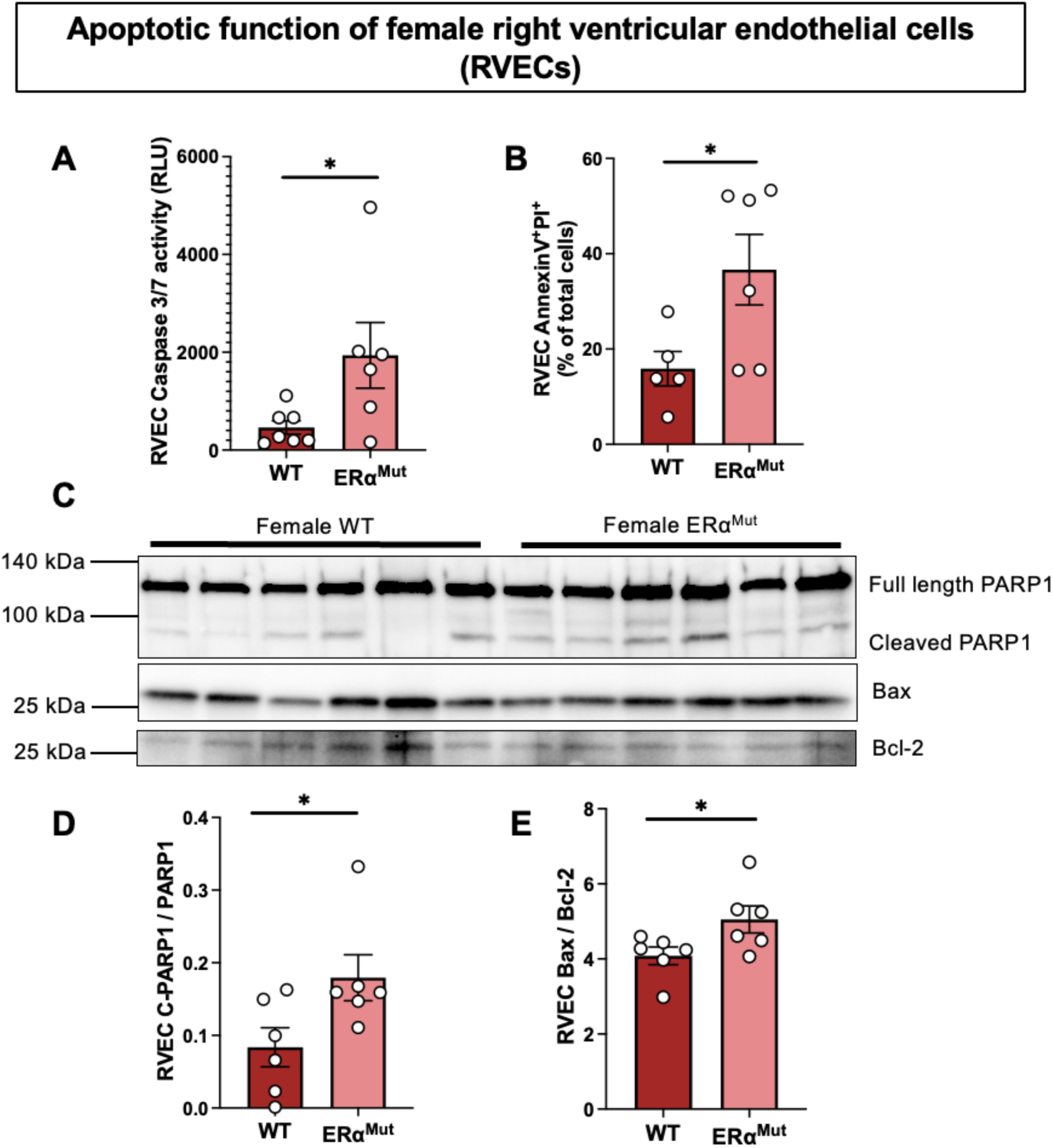
Loss of ERα exerts pro-apoptotic effects in female RVECs. (**A**) Caspase 3/7 activity assay in female WT and ERα^Mut^ RVECs grown for 24h. (**B**) Quantification of Annexin V^+^ and Propidium Iodide (PI)^+^ population in female WT and ERα^Mut^ RVECs grown for 24h. (**C**) Representative images of Western blots and quantification of cleaved/full length PARP1 (**D**) and Bax/Bcl-2 (**E**) grown for 24h. Each data point = one animal. * p<0.05, ** p<0.01; *** p<0.001 by ANOVA with Tukey’s post-hoc analysis. Error bars represent mean ± SEM. Error bars represent mean ± SEM. Scale bars, 50µm

To identify pathways and genes regulated by ERα in RVECs, angiogenesis PCR array was performed on female and male ERα^Mut^ versus WT RVECs (**Supplementary Fig. 3A**). The top 5 ranked differentially expressed genes (DEGs; including up-regulated and down-regulated) from each group are listed in **Supplementary Fig. 3B**. We then compared genes that were altered in female and male WT vs ERα^Mut^. Gene Ontology Biological Process analysis revealed that cell migration was the process with the most altered DEGs in both male (identified as cell migration, **Supplementary Fig. 2J**) and female RVECs (identified as taxis, **Fig. 2J**). Of note, leptin, a regulator of metabolism and angiogenesis^35–38^, was consistently downregulated with the loss of ERα (increased in WT), both in male (**Supplementary Fig. 2K**) and female RVECs (**Fig. 2K**).

To identify mechanisms underlying reduced angiogenic function in female ERα^Mut^ RVECs, we assessed apoptosis-related endpoints. Female ERα^Mut^ RVECs exhibited higher caspase 3/7 activity (**Fig. 2A**) and apoptotic cell percentages by AnnexinV-PI staining (**Fig. 2B**). After serum starvation, female ERα^Mut^ RVECs also exhibited higher cleaved PARP1 levels (Fig. **2C,D**) and higher Bax/Bcl-2 ratios (Fig. **2C,E**) than WT, suggesting increased pro-apoptotic signaling with loss of functioning ERα. Together, the data in Fig. 1&2 indicate that loss of functioning ERα in female RVECs results in increased propensity for apoptosis but also increased proliferation, with the latter potentially being a compensatory response for the increased cell death. These data also identify ERα as a sexually dimorphic pro-angiogenic mediator in RVECs.

### ERα protects RV capillaries from rarefaction in female MCT-PH rats

We next studied whether ERα regulates RVEC angiogenesis to prevent RV capillary rarefaction. Male and female WT and ERα^Mut^ rats were injected with MCT^15, 39, 40^ (**Fig. 3A**). As expected, and consistent with prior data, female WT rats exhibited only minimal increases in RVSP or Fulton index (**Fig. 3B,C**)^41–43^. On the other hand, as reported previously, female ERα^Mut^ rats exhibited a doubling in RVSP and Fulton index (**Fig. 3B,C**), suggesting ERα is protective against RVSP increase and RVH in females. Male WT MCT rats, on the other hand, exhibited expected increases in RVSP and Fulton index after MCT, but these changes were not affected by loss of functioning ERα (**Supplementary Fig. 4B,C**).

**Figure 3.**
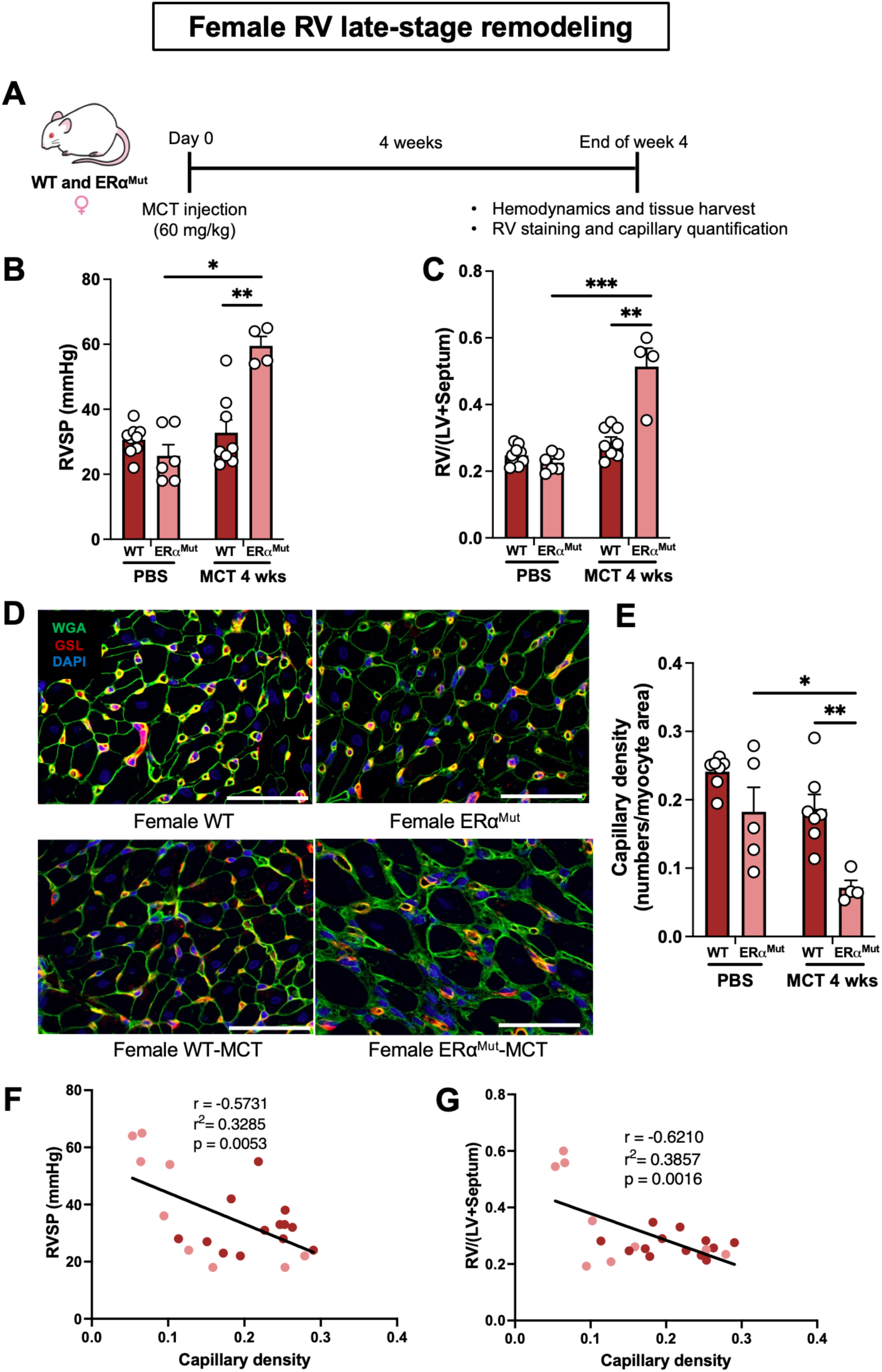
ERα deficiency results in PH and decreased RV capillary density in female MCT rats. (**A**) Schematic of experimental design. (**B-E**) Effects of ERα in female rats on RV systolic pressure (RVSP), RV hypertrophy and RV capillary density. (**B**) RVSP and (**C**) Fulton index in female WT and ERα^Mut^ rats treated with MCT or phosphate buffered saline (PBS; vehicle control). (**D**) Representative images of RV lectin staining and (**E**) Quantification of vascularization (as capillary per myocyte area) in female WT and ERα^Mut^ rats. Pearson correlation of vascularization (capillary per myocyte area) and (**F**) RVSP or (**G**) Fulton index. Dark dots indicate WT rats and light dots indicate ERα^Mut^ rats. Each data point represents one animal. WGA (Wheat Germ Agglutinin), cell membrane. GSL (Griffonia Simplicifolia Lectin), endothelial cells. Each data point = one animal. * p<0.05, ** p<0.01; *** p<0.001 by ANOVA with Tukey’s post-hoc analysis. Error bars represent mean ± SEM. Error bars represent mean ± SEM. Scale bars, 50µm

Next, we investigated RV capillary density in female PH rats after 4 weeks of MCT (**Fig. 3D,E**). Lectin staining revealed that female WT rats are resistant to MCT-induced RV capillary loss (**Fig. 3E**). Accompanying increases in RVSP and Fulton index, loss of ERα caused severe (∼65%) RV capillary rarefaction (**Fig. 3E**). Both RVSP (**Fig. 3F**) and RVH (**Fig. 3G**) demonstrated an inverse correlation with RV capillary density per myocyte area, suggesting a potential role for sufficient RV capillary density in preventing RVH. Taken together, these data suggest ERα exerts protective and pro-angiogenic effects in the female RV.

Interestingly, we did not observe significant RV capillary loss in male WT PH rats, and only male ERα^Mut^ PH rats exhibited RV capillary rarefaction (**Supplementary Fig. 4D,E**). Unlike female rats, RV capillary density per myocyte area only inversely correlated with Fulton index (**Supplementary Fig. 4G**), but not RVSP (**Supplementary Fig. 4F**).

### ERα protects female rats from early onset of RVH and RV capillary rarefaction after MCT exposure

To understand how ERα regulates RV capillary density in early stages of PH-induced RVH, we performed a time course experiment in male and female WT and ERα^Mut^ MCT rats. RVs were harvested at day 3 and day 10 after MCT injection. Interestingly, 10 days after MCT injection, Fulton index as well as RV cardiomyocyte cross-sectional area were significantly increased in female MCT ERα^Mut^ but not WT rats (**Fig. 4B,C**). This was accompanied by a significant decrease in RV capillary density (**Fig. 4D-G**). In contrast, there was no RVH and only a trend for RV capillary loss in male rats, with no difference between WT and ERα^Mut^ (**Supplementary Fig. 5B-G**). These data highlight the importance of ERα’s protective and pro-angiogenic role in the female RV even in early stages of PH and RV remodeling.

**Figure 4.**
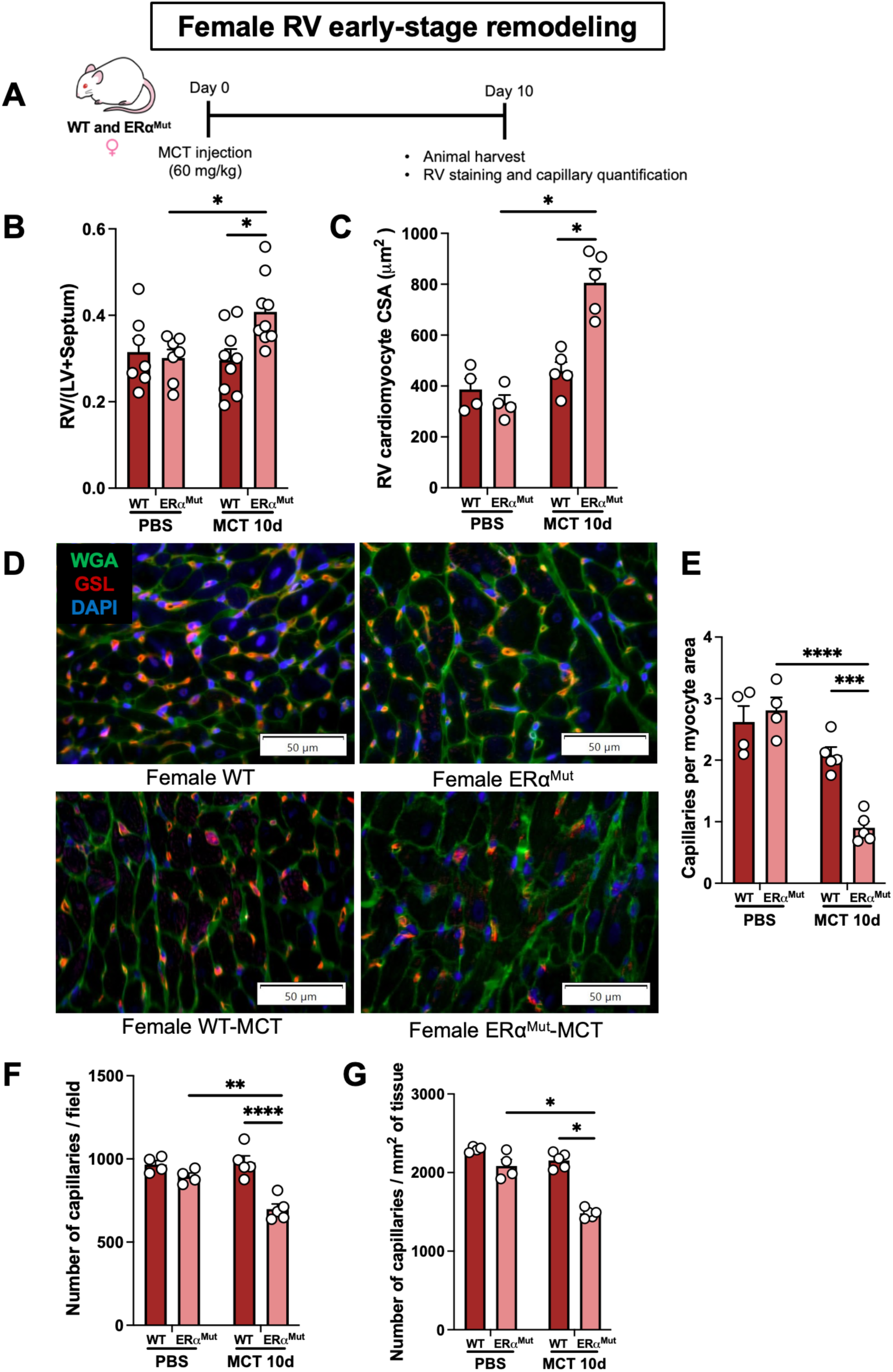
Loss of functional ERα results in early RV capillary rarefaction after MCT injection in female rats. (**A**) Schematic of experimental design. (**B-G**) Effects of ERα on RV hypertrophy and RV capillary density in female rats treated with MCT or PBS. (**B**) Fulton index and (**C**) RV cardiomyocyte size in female WT and ERα^Mut^ MCT-PH rats. (**D**) Representative images and (**E-G**) quantification of RV capillary density via lectin staining. WGA (Wheat Germ Agglutinin), cell membrane. GSL (Griffonia Simplicifolia Lectin), endothelial cells. CSA, cross-sectional area. Each data point = one animal. * p<0.05, ** p<0.01; *** p<0.001; **** p<0.0001 by ANOVA with Tukey’s post-hoc analysis. Error bars represent mean ± SEM. Error bars represent mean ± SEM. Scale bars, 50µm

**Figure 5.**
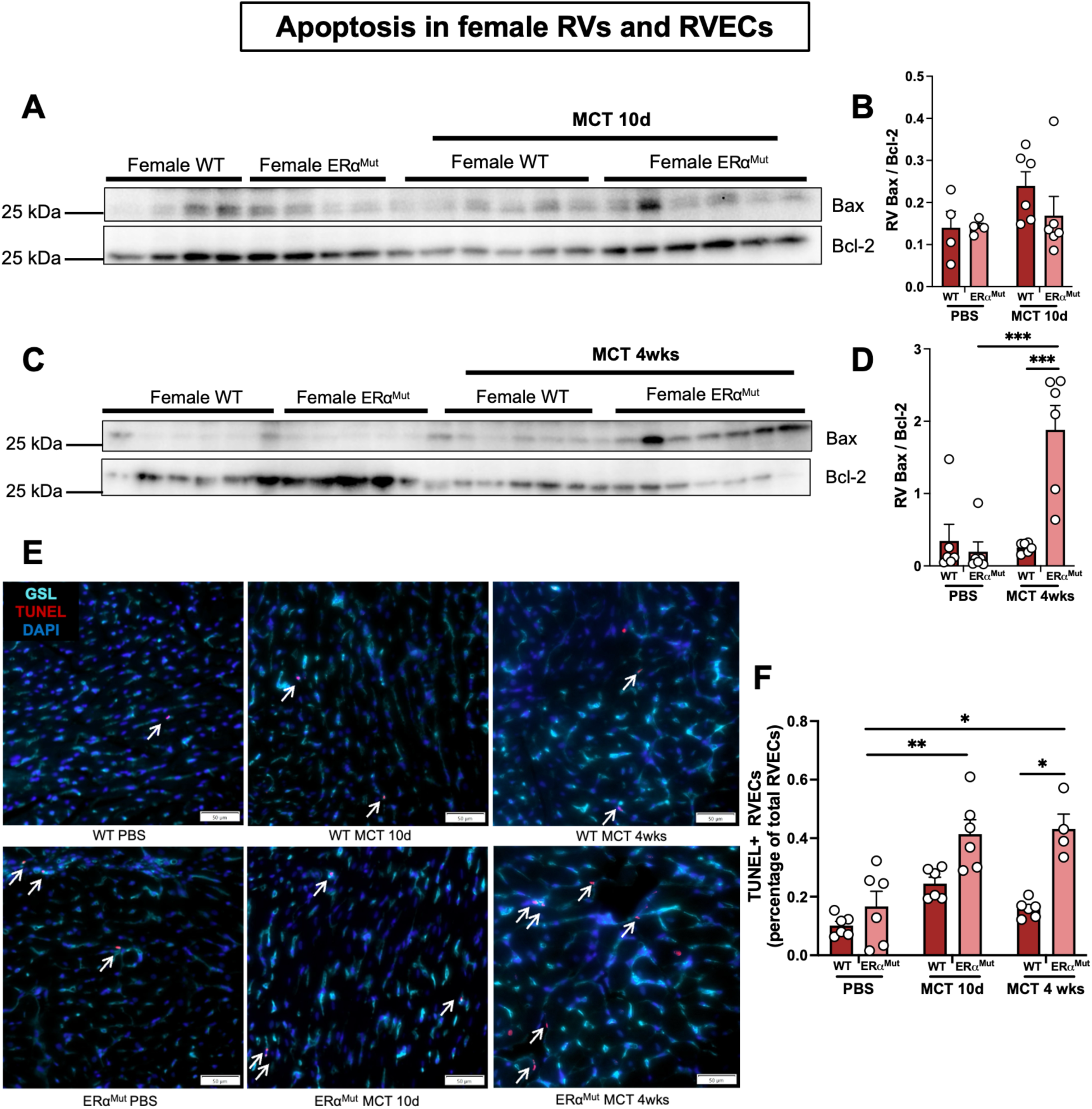
Female ERα^Mut^ rats exhibit increased RVEC apoptosis after MCT treatment. **(A-D)** Representative images and quantification of Western blot analysis of Bax and Bcl-2 in female WT PBS or ERα^Mut^ RV tissues ten days and four weeks after MCT administration, respectively. (**E**) Representative images of TUNEL^+^ RVECs (RVECs stained with griffonia simplicifolia lectin; GSL). (**F**) Quantification of TUNEL^+^ RVECs (as percentage of total RVEC number). White arrows, TUNEL^+^ RVECs. PBS, phosphate buffered saline. Each data point represents one animal. Each data point = one animal. * p<0.05, ** p<0.01; *** p<0.001 by ANOVA with Tukey’s post-hoc analysis. Error bars represent mean ± SEM. Error bars represent mean ± SEM. Scale bars, 50µm

Capillary staining of male and female RVs three days after MCT administration showed similar capillary density as compared to controls and did not reveal a sex bias or ERα regulation (**Supplementary Fig. 6D-G & 7D-G**).

### Female ERα^Mut^ rats exhibit more severe pulmonary vascular remodeling during early stages of PH development

In PH, RVH is induced by increased RV afterload from pulmonary vascular remodeling. We recently reported greater PA muscularization in female ERα^Mut^ rats^44^. Therefore, we examined if the RVH and RV capillary rarefaction noted in female ERα^Mut^ rats at 10 days are associated with more severe pulmonary vascular remodeling. Indeed, female ERα^Mut^ rats demonstrated fewer partially muscularized vessels and more fully muscularized vessels (**Supplementary Fig. 8A,B**). These data indicate that the RVH and RV capillary loss in female ERα^Mut^ rats 10 days after MCT may be due to enhanced pulmonary vascular remodeling. On the other hand, we observed a modest increase in the number of partially muscularized vessels in male ERα^Mut^ rats with MCT treatment at day 10, but no difference in nonmuscularized or fully muscularized PAs (**Supplementary Fig. 9A,B**). These data mirror the RV capillary density data, where female ERα^Mut^ rats exhibit more RVH and RV capillary rarefaction than WT, paralleled by more severe pulmonary vascular remodeling. Male ERα^Mut^ rats on the other hand exhibit no increase in RVH, RV capillarization, and pulmonary vascular remodeling as compared to WT (**Supplementary Fig. 5B-G**).

### ERα^Mut^ inhibits apoptotic signaling in RVECs from female MCT rats

We next examined whether enhanced RVEC apoptosis contributes to the reduced angiogenic capacity and capillary rarefaction observed in female ERα^Mut^ rats at early and late timepoints. Bax/Bcl-2 ratio was not different in the RV of female WT and ERα^Mut^ rats 10 days after MCT (**Fig. 7A,B**). However, at 4 weeks after MCT treatment, female ERα^Mut^ rats demonstrated a higher Bax/Bcl-2 ratio (**Fig. 7C,D**) compared to WT. TUNEL staining revealed a greater number of TUNEL⁺ RVECs in female ERα^Mut^ rats at both 10 days and 4 weeks post-MCT treatment relative to WT rats (**Fig. 7E,F**). Together, these results suggest that functional ERα protects the female RV from endothelial apoptotic cell death during both early and late stages of MCT-induced remodeling.

### ERα regulates migration and apopotosis pathways in female capillary and endocardial RVECs

To identify genes and pathways regulated by ERα in female RVECs, we performed snRNA-seq on RV tissues from female WT and ERα^Mut^ rats at both 10 days and 4 weeks after MCT administration (**Fig. 6A**). We discovered five distinct populations of RVECs: arterial ECs, capillary ECs, endocardial ECs, lymphatic ECs, and venous ECs (**Fig. 6B,C**). Featured genes of each RVEC sub-population are shown in **Fig. 6B**. At 4 weeks, there was a reduction in the endocardial EC population and an increase in the capillary EC population in female ERα^Mut^ (**Fig. 6D,E**). We identified higher numbers of DEGs between female WT and ERα^Mut^ across all 5 sub-populations of RVECs at 10 days compared to 4 weeks after MCT (**Fig. 7A,B**). This suggests ERα regulates gene expression predominantly at early stages of RVH. In addition, Qiagen Ingenuity® Pathway Analysis uncovered differentially regulated pathways between WT and ERα^Mut^ (**Fig. 7C-F**). At 10 days, migration was higher in female WT capillary and endocardial RVECs than in ERα^Mut^ cells (**Fig. 7C,E**), suggesting that loss of ERα reduces migration in vivo. Moreover, compared to ERα^Mut^ cells, female WT endocardial RVECs demonstrated increased cell survival and a less apoptotic phenotype (**Fig. 7E**), linking loss of functioning ERα to increased apoptotic cell death. At 4 weeks, female WT capillary RVECs displayed increased survival programming compared to ERα^Mut^ cells (**Fig. 7D**), again linking loss of functioning ERα to increased cell death. Other significantly increased pathways in WT included organization of cytoskeleton, microtubule dynamics, and morphogenesis of cellular protrusions. Taken together, snRNA-seq data suggest that ERα plays a fundamental role in maintaining cell viability and RVEC migration during the development of RV failure.

**Figure 6.**
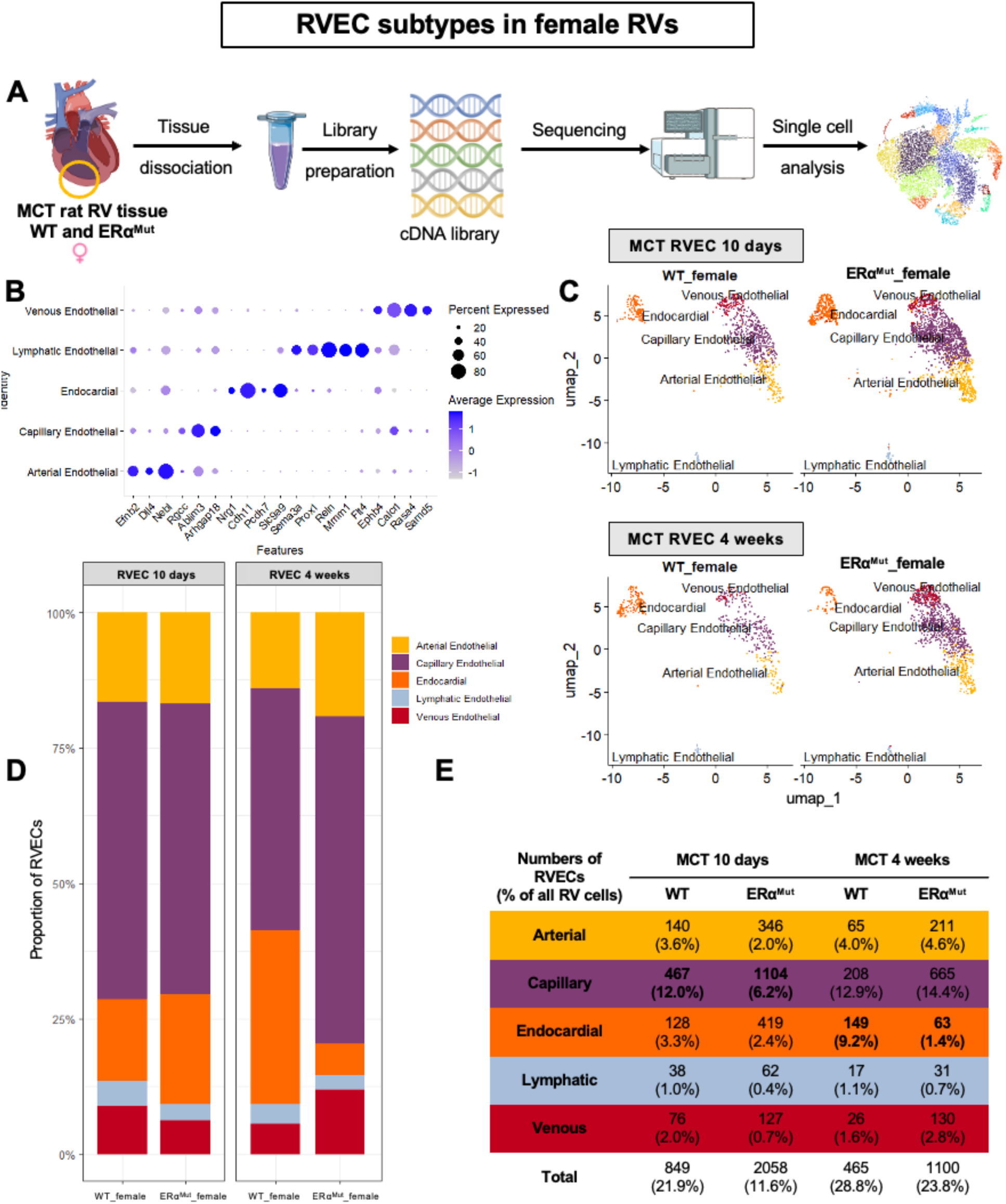
Single nucleus RNA-sequencing of female MCT-PH RVs reveals five distinct populations of RVECs. (**A**) Schematic of snRNA-Seq experimental design. (**B**) Featured gene expressions of each RVEC sub-population. (**C**) Uniform Manifold Approximation and Projection (UMAP) plots of RVEC sub-populations at ten days and four weeks.(**D**) Cellular proportion plot of RVEC sub-populations. (**E**) Total numbers and percentages of RVEC sub-populations in relation to total RV cell number (including all RV cell types). n=2 animals per group.

**Figure 7.**
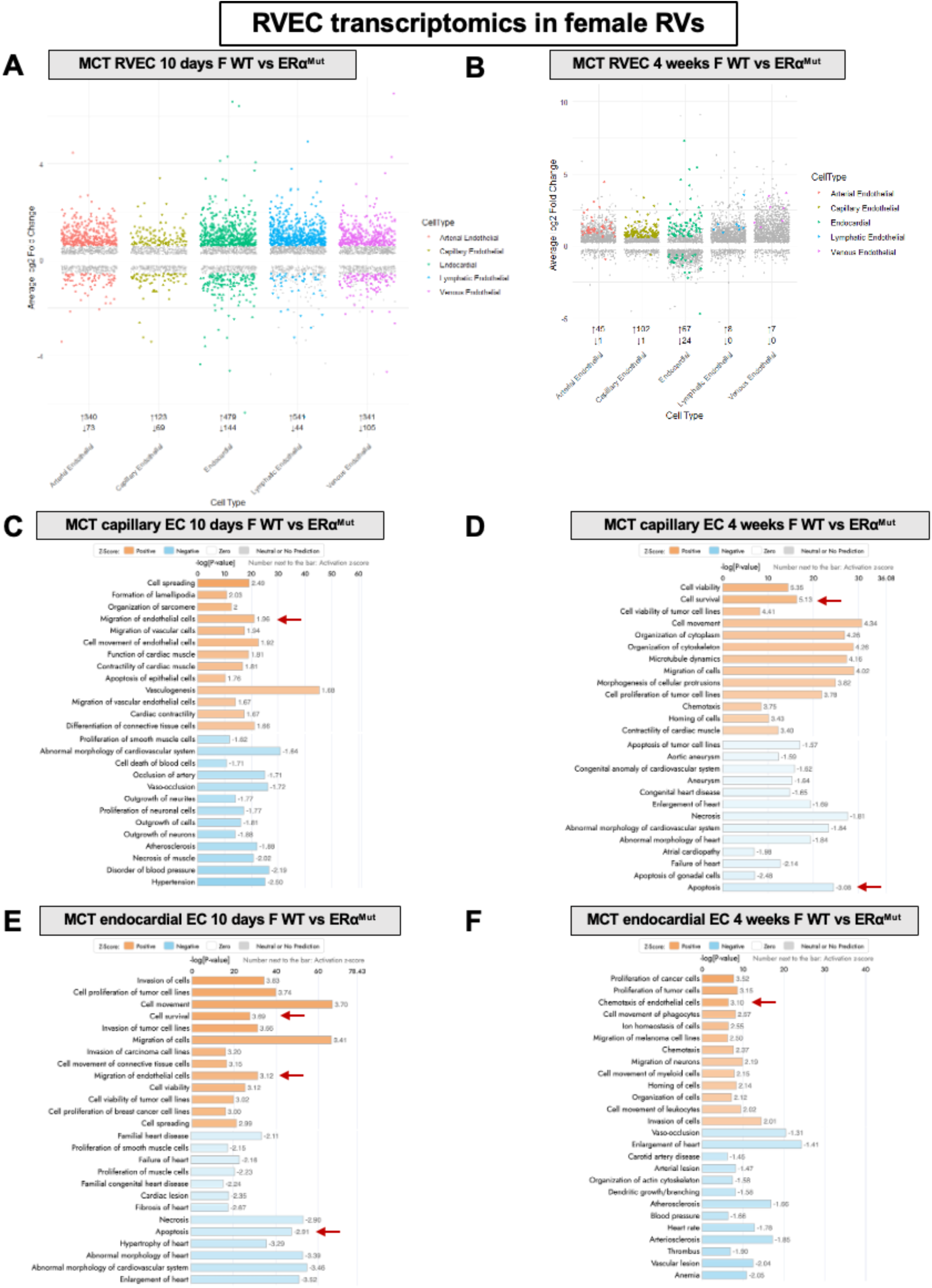
ERα regulates early changes in RVEC gene programs in female MCT-PH RVs. **(A, B)** Differentially expressed genes (DEGs) of female WT ERα^Mut^ RVECs ten days (**A**) and four weeks (**B**) after MCT administration. **(C-F)** Qiagen IPA disease and function pathway analysis in female capillary and endocardial RVECs in WT vs ERα^Mut^ cells at ten days (**C, D**) and four weeks (**E,F**) after MCT administration. Pathways are ranked by Z-score. Only top upregulated and downregulated pathways are shown.

## Discussion

We report for the first time that ERα regulates RVEC function and RV angiogenesis in a sexually dimorphic manner. Loss of functioning ERα resulted in reduced RVEC pseudovascular network formation and migration in male and female RVECs at baseline conditions, as well as increased apoptosis in female RVECs. In vivo, loss of functioning ERα resulted in reduced RV capillary density in female MCT-PH rats. In addition, we observed early onset of RVH and capillary loss with the loss of ERα in females. These findings were accompanied by a higher number of apoptotic RVECs in female ERα^Mut^ MCT-PH RVs. snRNA-Seq in female MCT-PH RVs revealed previously undescribed RVEC subtypes that are under transcriptional regulation of ERα. In particular, downregulation of migration and upregulation of apoptosis pathways were identified in female ERα^Mut^ MCT-PH rats. Together, these data indicate that ERα’s effects on RVEC function and RV angiogenesis exhibit a female sex bias.

In response to the increased pressure overload caused by PH, the RV adapts via cardiomyocyte hypertrophy and increased cardiac contractility^6, 7^. In the initial adaptive stage, there is increased capillary density and vascularization^45^. This is achieved by up-regulating pro-angiogenic signaling^46^ and RVEC proliferation^45^. The molecular switch that drives the transition from adaptive vascular maintenance to maladaptive capillary loss, however, remains poorly understood. Our findings implicate ERα as a novel regulator of RVEC function and RV vascularization during PH progression.

We previously demonstrated beneficial effects of ERα on RV cardiomyocytes^16^. The current study expands those findings by investigating the role of ERα in RVECs (another cell type central to the pathogenesis of RVF^13, 47^). This was motivated by the established role of RVEC dysfunction in the pathogenesis of RVF, the known sexual dimorphisms in RV adaptation to PH, and the well-characterized vascular-protective effects of ERα in systemic and left ventricular endothelium. Vascular ECs are known to express all three types of estrogen receptors (ERα, ERβ and GPER), and the function of ERα in the vasculature has been well studied. ERα has EC-protective effects in various types of vascular injury^48, 49^. For example, ERα is directly linked to eNOS expression^50, 51^ and regulation of vascular tone^52^. Mouse models with ERα deletion and/or mutations have delineated the function of ERα in the vasculature (reviewed in ^53^). For example, ERα is essential for the beneficial effects of E2 on re-endothelialization and injury-induced medial hyperplasia^54^. Expression of ERα in vascular ECs is modulated by estrogen status and strongly related to endothelial-dependent dilation in women^55^. Variations in *ESR1* (the gene encoding ERα) have been linked to myocardial infarction in both men and postmenopausal women^56, 57^, highlighting the protective effect of ERα in the cardiovascular system.

ERα promotes EC migration through the c-Src and focal adhesion kinase pathway^58^, as well as the RhoA/Moesin pathway^59^. Our findings expand this observation and, for the first time, demonstrate stimulatory effects of ERα on migration in RVECs. Effects of ERα on apoptosis are less understood. E2 inhibits TNF-α induced EC apoptosis^60^ and ERβ protects human umbilical vein ECs against apoptosis^61^. We, for the first time, discovered that loss of ERα causes increased apoptosis in female RVECs, suggesting that ERα-mediated suppression of endothelial apoptosis contributes to the maintenance of RV capillary density in females. This is particularly relevant given clinical evidence showing increased apoptotic vessels in RVs from female PAH patients compared to control RVs^62^. Upregulating RV angiogenesis, inhibiting RVEC apoptosis, or increasing RVEC migration are potential new mechanisms by which ERα maintains RV function. Further research is needed to uncover the downstream pathways that are affected by the loss of ERα.

Our research also unveiled a sexual dimorphic role of ERα in protecting RV from failure. In males MCT-PH rats, RVSP elevations, Fulton indices, and capillary density reductions are comparable between WT and ERα^Mut^, suggesting that ERα does not provide protective benefits in the male RV and is playing a dispensable role in RVH induced by PH. On the other hand, femaleMCT-PH rats are relying upon ERα for resilience against PH-induced RVH and vascularization defects.

Previous studies identified sex differences in the timing and severity of PH-induced RV dysfunction^63, 64^. In male WT rats, significant elevation of mPAP and Fulton index were observed after 2 weeks of MCT injection^64^, as well as reduced RV capillary density^63^. Female rats, on the other hand, demonstrated less pronounced RVH and dilatation^63^. This indicates that female WT rats are more resistant to RVH development caused by MCT at early stages. Given the beneficial effects provided by ERα in the vasculature, we focused on ERα’s pro-angiogenic effects in RV. Our time course experiments suggest that 1) RV remodeling in female rats happens earlier post-MCT than in male rats; 2) RV capillary density in female rats contributes more to RVH and RV dysfunction than in their male counterparts; and 3) ERα is a critical mediator of RV protection and capillary maintenance in females during PH development.

The ERα loss-of-function rat model investigated in this study was generated through CRISPR/Cas9 as previously described^16^. Our rats exhibit a host of phenotypes in the pulmonary vasculature and RV when subjected to experimental PH or RV pressure overload. Interestingly, compared to male ERα^Mut^ rats, female ERα^Mut^ rats exhibit a more profound phenotype. When subjected to MCT-PH, female ERα^Mut^ rats exhibit higher RVSP and Fulton index, more pulmonary vascular remodeling and more profound decreases in RV function, whereas male WT and ERα^Mut^ rats have comparable disease phenotypes^44^. At a cellular level, we discovered a novel E2-ERα-BMPR2-apelin axis in RV cardiomyocytes. In light of the focus of our paper, this is of particular interest, since both BMPR2 as well as apelin are known pro-angiogenic mediators^16^. We also found that the RV cardiomyocyte secretome increases RVEC angiogenic ability in vitro, and that the pro-angiogenic function of RV cardiomyocytes is blunted when apelin is silenced^16^. However, in our snRNA-Seq analyses in female MCT RVECs we did not detect BMPR2 nor apelin being differentially expressed between WT and ERα^Mut^. This may be a reflection of this axis being suppressed in the setting of PH and RV failure or indicate that BMPR2 or apelin are not under transcriptional control of ERα in RVECs.

Our snRNA-Seq experiments, for the first time, identified five distinct sub-populations of RVECs. This allows us to better understand RVEC dynamics and functions during the development of RVH. We noticed capillary and endocardial RVECs were the most changed populations among the five subpopulations. Of note, ventricular endocardial cells play an vital role in coronary artery angiogenesis as these cells have the potential to differentiate into arterial or capillary endothelial cells^65^. However, the transition from endocardial to capillary ECs in the context of RVH remains elusive. Not surprisingly, ERα differentially regulates the gene programming of RVEC sub-populations during the development of RVH. The observation that capillary and endocardial RVEC numbers are drastically changed suggests that ERα provides beneficial effects by regulating these cells. The drastic decline of endocardial ECs at 4 weeks after MCT treatment points to a new mechanism of angiogenesis pertubation in RV failure where a loss of endocardial EC could potentially explain the insufficient RV angiogenesis. Maintaining endocardial EC viability could therefore be a novel therapeutic avenue. Furthermore, the DEG analysis indicates that transcriptome changes regulated by ERα happen early in the disease progression. This suggests a need for early intervention and identifies enhancing ERα signaling as a potential novel therapeutic strategy to protect the RV in PH.

One limitation of our research is the use of a two-diemsional method for quantifying the three-dimensional capillary structure of the RV. Stereology is considered the gold standard for assessing three-dimensional structures. We did not consistently use stereology due to its limitation of requiring the entire RV to be fixed, which would not allow for isolating RVECs or performing biochemical analyses. Several important prior studies in the field employed two-dimensional analysis methods for quantifying RV vascularization^12, 13, 66, 67^. In addition, we and others^13^ demonstrated that findings of decreased vascular density in vivo are accompanied by decreased pseudo-vascular network formation and impaired angiogenic behavior in cultured RVECs, thus supporting the in vivo findings. A recent paper suggests that it may not be the loss of capillaries in RV, but rather, a lack of sufficient contact area between RVECs and RV cardiomyocytes that causes maladaptive RVH^17^.

We focused on studying ERα without its ligand E2. This is due to the fact that ERα can elicit downstream signaling even in absence of activation by E2^53^. It has been shown that there is increased susceptibility to PH in women with premature menopause^68^ and we found reduced RVEC and RV cardiomyocyte expression levels of ERα in patients with RV failure^16^. This suggests that reduced ERα activity could be a contributor to RV failure. Therefore, we chose to study ERα without the influence of E2 as a first step. Studies employing E2 supplementation are currently ongoing in our laboratory.

In summary, our research provides novel insights into how ERα regulates RV angiogenesis and RVEC function and identifies ERα as being essential for preventing RV failure and RV capillary rarefaction in females with PH. Our reseach also sheds light on the sexually dimorphic role of ERα in males and females with PH.

## Supporting information

All supplementary information

## Acknowledgement

We thank the Genome Technology Access Center at the McDonnell Genome Institute at Washington University School of Medicine for help with genomic analysis. The Center is partially supported by NCI Cancer Center Support Grant #P30 CA91842 to the Siteman Cancer Center from the National Center for Research Resources (NCRR). This publication is solely the responsibility of the authors and does not necessarily represent the official view of NCRR or NIH.

## References

1. Mocumbi A, Humbert M, Saxena A, Jing ZC, Sliwa K, Thienemann F, Archer SL, Stewart S. Pulmonary hypertension. Nat Rev Dis Primers 2024;10:1.

2. Vonk Noordegraaf A, Galiè N. The role of the right ventricle in pulmonary arterial hypertension. European Respiratory Review 2011;20:243–253.

3. van de Veerdonk MC, Kind T, Marcus JT, Mauritz GJ, Heymans MW, Bogaard HJ, Boonstra A, Marques KM, Westerhof N, Vonk-Noordegraaf A. Progressive right ventricular dysfunction in patients with pulmonary arterial hypertension responding to therapy. J Am Coll Cardiol 2011;58:2511–2519.

4. Ryan JJ, Huston J, Kutty S, Hatton ND, Bowman L, Tian L, Herr JE, Johri AM, Archer SL. Right ventricular adaptation and failure in pulmonary arterial hypertension. Can J Cardiol 2015;31:391–406.

5. Padang R, Chandrashekar N, Indrabhinduwat M, Scott CG, Luis SA, Chandrasekaran K, Michelena HI, Nkomo VT, Pislaru SV, Pellikka PA, Kane GC. Aetiology and outcomes of severe right ventricular dysfunction. European Heart Journal 2020;41:1273–1282.

6. Vonk Noordegraaf A, Westerhof BE, Westerhof N. The Relationship Between the Right Ventricle and its Load in Pulmonary Hypertension. J Am Coll Cardiol 2017;69:236–243.

7. Vonk Noordegraaf A, Chin KM, Haddad F, Hassoun PM, Hemnes AR, Hopkins SR, Kawut SM, Langleben D, Lumens J, Naeije R. Pathophysiology of the right ventricle and of the pulmonary circulation in pulmonary hypertension: an update. Eur Respir J 2019;53.

8. Houston BA, Brittain EL, Tedford RJ. Right Ventricular Failure. New England Journal of Medicine 2023;388:1111–1125.

9. Zelt JGE, Chaudhary KR, Cadete VJ, Mielniczuk LM, Stewart DJ. Medical Therapy for Heart Failure Associated With Pulmonary Hypertension. Circulation Research 2019;124:1551–1567.

10. Evans CE, Cober ND, Dai Z, Stewart DJ, Zhao YY. Endothelial cells in the pathogenesis of pulmonary arterial hypertension. Eur Respir J 2021;58.

11. Cober ND, VandenBroek MM, Ormiston ML, Stewart DJ. Evolving Concepts in Endothelial Pathobiology of Pulmonary Arterial Hypertension. Hypertension 2022;79:1580–1590.

12. Piao L, Fang YH, Parikh K, Ryan JJ, Toth PT, Archer SL. Cardiac glutaminolysis: a maladaptive cancer metabolism pathway in the right ventricle in pulmonary hypertension. J Mol Med (Berl) 2013;91:1185–1197.

13. Potus F, Ruffenach G, Dahou A, Thebault C, Breuils-Bonnet S, Tremblay È, Nadeau V, Paradis R, Graydon C, Wong R, Johnson I, Paulin R, Lajoie AC, Perron J, Charbonneau E, Joubert P, Pibarot P, Michelakis ED, Provencher S, Bonnet S. Downregulation of MicroRNA-126 Contributes to the Failing Right Ventricle in Pulmonary Arterial Hypertension. Circulation 2015;132:932–943.

14. Graham BB, Kumar R, Mickael C, Kassa B, Koyanagi D, Sanders L, Zhang L, Perez M, Hernandez-Saavedra D, Valencia C, Dixon K, Harral J, Loomis Z, Irwin D, Nemkov T, D’Alessandro A, Stenmark KR, Tuder RM. Vascular Adaptation of the Right Ventricle in Experimental Pulmonary Hypertension. Am J Respir Cell Mol Biol 2018;59:479–489.

15. Frump AL, Bonnet S, de Jesus Perez VA, Lahm T. Emerging role of angiogenesis in adaptive and maladaptive right ventricular remodeling in pulmonary hypertension. Am J Physiol Lung Cell Mol Physiol 2018;314:L443–l460.

16. Frump AL, Albrecht M, Yakubov B, Breuils-Bonnet S, Nadeau V, Tremblay E, Potus F, Omura J, Cook T, Fisher A, Rodriguez B, Brown RD, Stenmark KR, Rubinstein CD, Krentz K, Tabima DM, Li R, Sun X, Chesler NC, Provencher S, Bonnet S, Lahm T. 17β-Estradiol and estrogen receptor α protect right ventricular function in pulmonary hypertension via BMPR2 and apelin. J Clin Invest 2021;131.

17. Ichimura K, Boehm M, Andruska AM, Zhang F, Schimmel K, Bonham S, Kabiri A, Kheyfets VO, Ichimura S, Reddy S, Mao Y, Zhang T, Wang GX, Santana EJ, Tian X, Essafri I, Vinh R, Tian W, Nicolls MR, Yajima S, Shudo Y, MacArthur JW, Woo YJ, Metzger RJ, Spiekerkoetter E. 3D Imaging Reveals Complex Microvascular Remodeling in the Right Ventricle in Pulmonary Hypertension. Circ Res 2024;135:60–75.

18. Tello K, Richter MJ, Yogeswaran A, Ghofrani HA, Naeije R, Vanderpool R, Gall H, Tedford RJ, Seeger W, Lahm T. Sex Differences in Right Ventricular-Pulmonary Arterial Coupling in Pulmonary Arterial Hypertension. Am J Respir Crit Care Med 2020;202:1042–1046.

19. Jacobs W, van de Veerdonk MC, Trip P, de Man F, Heymans MW, Marcus JT, Kawut SM, Bogaard HJ, Boonstra A, Vonk Noordegraaf A. The right ventricle explains sex differences in survival in idiopathic pulmonary arterial hypertension. Chest 2014;145:1230–1236.

20. Umar S, Iorga A, Matori H, Nadadur RD, Li J, Maltese F, van der Laarse A, Eghbali M. Estrogen rescues preexisting severe pulmonary hypertension in rats. Am J Respir Crit Care Med 2011;184:715–723.

21. Ventetuolo CE, Ouyang P, Bluemke DA, Tandri H, Barr RG, Bagiella E, Cappola AR, Bristow MR, Johnson C, Kronmal RA, Kizer JR, Lima JA, Kawut SM. Sex hormones are associated with right ventricular structure and function: The MESA-right ventricle study. Am J Respir Crit Care Med 2011;183:659–667.

22. Frump AL, Goss KN, Vayl A, Albrecht M, Fisher A, Tursunova R, Fierst J, Whitson J, Cucci AR, Brown MB, Lahm T. Estradiol improves right ventricular function in rats with severe angioproliferative pulmonary hypertension: effects of endogenous and exogenous sex hormones. Am J Physiol Lung Cell Mol Physiol 2015;308:L873–890.

23. Lahm T, Albrecht M, Fisher AJ, Selej M, Patel NG, Brown JA, Justice MJ, Brown MB, Van Demark M, Trulock KM, Dieudonne D, Reddy JG, Presson RG, Petrache I. 17β-Estradiol attenuates hypoxic pulmonary hypertension via estrogen receptor-mediated effects. Am J Respir Crit Care Med 2012;185:965–980.

24. Billon-Galés A, Fontaine C, Filipe C, Douin-Echinard V, Fouque M-J, Flouriot G, Gourdy P, Lenfant F, Laurell H, Krust A, Chambon P, Arnal J-F. The transactivating function 1 of estrogen receptor α is dispensable for the vasculoprotective actions of 17β-estradiol. Proceedings of the National Academy of Sciences 2009;106:2053–2058.

25. Clere N, Lauret E, Malthiery Y, Andriantsitohaina R, Faure S. Estrogen receptor alpha as a key target of organochlorines to promote angiogenesis. Angiogenesis 2012;15:745–760.

26. Mahmoodzadeh S, Leber J, Zhang X, Jaisser F, Messaoudi S, Morano I, Furth PA, Dworatzek E, Regitz-Zagrosek V. Cardiomyocyte-specific Estrogen Receptor Alpha Increases Angiogenesis, Lymphangiogenesis and Reduces Fibrosis in the Female Mouse Heart Post-Myocardial Infarction. J Cell Sci Ther 2014;5:153.

27. Lahm T, Douglas IS, Archer SL, Bogaard HJ, Chesler NC, Haddad F, Hemnes AR, Kawut SM, Kline JA, Kolb TM, Mathai SC, Mercier O, Michelakis ED, Naeije R, Tuder RM, Ventetuolo CE, Vieillard-Baron A, Voelkel NF, Vonk-Noordegraaf A, Hassoun PM. Assessment of Right Ventricular Function in the Research Setting: Knowledge Gaps and Pathways Forward. An Official American Thoracic Society Research Statement. Am J Respir Crit Care Med 2018;198:e15–e43.

28. Provencher S, Archer SL, Ramirez FD, Hibbert B, Paulin R, Boucherat O, Lacasse Y, Bonnet S. Standards and Methodological Rigor in Pulmonary Arterial Hypertension Preclinical and Translational Research. Circ Res 2018;122:1021–1032.

29. Litviňuková M, Talavera-López C, Maatz H, Reichart D, Worth CL, Lindberg EL, Kanda M, Polanski K, Heinig M, Lee M, Nadelmann ER, Roberts K, Tuck L, Fasouli ES, DeLaughter DM, McDonough B, Wakimoto H, Gorham JM, Samari S, Mahbubani KT, Saeb-Parsy K, Patone G, Boyle JJ, Zhang H, Zhang H, Viveiros A, Oudit GY, Bayraktar OA, Seidman JG, Seidman CE, Noseda M, Hubner N, Teichmann SA. Cells of the adult human heart. Nature 2020;588:466–472.

30. Hocker JD, Poirion OB, Zhu F, Buchanan J, Zhang K, Chiou J, Wang T-M, Zhang Q, Hou X, Li YE, Zhang Y, Farah EN, Wang A, McCulloch AD, Gaulton KJ, Ren B, Chi NC, Preissl S. Cardiac cell type–specific gene regulatory programs and disease risk association. Science Advances 2021;7:eabf1444.

31. Koenig AL, Shchukina I, Amrute J, Andhey PS, Zaitsev K, Lai L, Bajpai G, Bredemeyer A, Smith G, Jones C, Terrebonne E, Rentschler SL, Artyomov MN, Lavine KJ. Single-cell transcriptomics reveals cell-type-specific diversification in human heart failure. Nature Cardiovascular Research 2022;1:263–280.

32. Ferrara N, Gerber HP, LeCouter J. The biology of VEGF and its receptors. Nat Med 2003;9:669–676.

33. Olsson AK, Dimberg A, Kreuger J, Claesson-Welsh L. VEGF receptor signalling -in control of vascular function. Nat Rev Mol Cell Biol 2006;7:359–371.

34. Lamalice L, Le Boeuf F, Huot J. Endothelial Cell Migration During Angiogenesis. Circulation Research 2007;100:782–794.

35. Bouloumié A, Drexler HCA, Lafontan M, Busse R. Leptin, the Product of Ob Gene, Promotes Angiogenesis. Circulation Research 1998;83:1059–1066.

36. Sierra-Honigmann MRo, Nath AK, Murakami C, García-Cardeña G, Papapetropoulos A, Sessa WC, Madge LA, Schechner JS, Schwabb MB, Polverini PJ, Flores-Riveros JR. Biological Action of Leptin as an Angiogenic Factor. Science 1998;281:1683–1686.

37. Cao R, Brakenhielm E, Wahlestedt C, Thyberg J, Cao Y. Leptin induces vascular permeability and synergistically stimulates angiogenesis with FGF-2 and VEGF. Proceedings of the National Academy of Sciences 2001;98:6390–6395.

38. Garonna E, Botham KM, Birdsey GM, Randi AM, Gonzalez-Perez RR, Wheeler-Jones CPD. Vascular Endothelial Growth Factor Receptor-2 Couples Cyclo-Oxygenase-2 with Pro-Angiogenic Actions of Leptin on Human Endothelial Cells. PLOS ONE 2011;6:e18823.

39. Stenmark KR, Meyrick B, Galie N, Mooi WJ, McMurtry IF. Animal models of pulmonary arterial hypertension: the hope for etiological discovery and pharmacological cure. American Journal of Physiology-Lung Cellular and Molecular Physiology 2009;297:L1013–L1032.

40. Boucherat O, Agrawal V, Lawrie A, Bonnet S. The Latest in Animal Models of Pulmonary Hypertension and Right Ventricular Failure. Circulation Research 2022;130:1466–1486.

41. Rafikova O, James J, Eccles CA, Kurdyukov S, Niihori M, Varghese MV, Rafikov R. Early progression of pulmonary hypertension in the monocrotaline model in males is associated with increased lung permeability. Biology of Sex Differences 2020;11:11.

42. Lookin O, Kuznetsov D, Protsenko Y. Sex differences in stretch-dependent effects on tension and Ca2+ transient of rat trabeculae in monocrotaline pulmonary hypertension. The Journal of Physiological Sciences 2015;65:89–98.

43. Goldenthal EI, D AW, Lynch JF. HORMONAL MODIFICATION OF THE SEX DIFFERENCES FOLLOWING MONOCROTALINE ADMINISTRATION. Toxicol Appl Pharmacol 1964;6:434–441.

44. Frump AL, Yakubov B, Walts A, Fisher A, Cook T, Chesler NC, Lahm T. Estrogen Receptor-α Exerts Endothelium-Protective Effects and Attenuates Pulmonary Hypertension. Am J Respir Cell Mol Biol 2023;68:341–344.

45. Kolb TM, Peabody J, Baddoura P, Fallica J, Mock JR, Singer BD, D’Alessio FR, Damarla M, Damico RL, Hassoun PM. Right Ventricular Angiogenesis is an Early Adaptive Response to Chronic Hypoxia-Induced Pulmonary Hypertension. Microcirculation 2015;22:724–736.

46. Partovian C, Adnot S, Eddahibi S, Teiger E, Levame M, Dreyfus P, Raffestin B, Frelin C. Heart and lung VEGF mRNA expression in rats with monocrotaline- or hypoxia-induced pulmonary hypertension. Am J Physiol 1998;275:H1948–1956.

47. Frump AL, Bonnet S, de Jesus Perez VA, Lahm T. Emerging role of angiogenesis in adaptive and maladaptive right ventricular remodeling in pulmonary hypertension. American Journal of Physiology-Lung Cellular and Molecular Physiology 2018;314:L443–L460.

48. Pare G, Krust A, Karas RH, Dupont S, Aronovitz M, Chambon P, Mendelsohn ME. Estrogen Receptor-α Mediates the Protective Effects of Estrogen Against Vascular Injury. Circulation Research 2002;90:1087–1092.

49. Liu P-Y, Fukuma N, Hiroi Y, Kunita A, Tokiwa H, Ueda K, Kariya T, Numata G, Adachi Y, Tajima M, Toyoda M, Li Y, Noma K, Harada M, Toko H, Ushiku T, Kanai Y, Takimoto E, Liao JK, Komuro I. Tie2-Cre–Induced Inactivation of Non-Nuclear Estrogen Receptor-α Signaling Abrogates Estrogen Protection Against Vascular Injury. JACC: Basic to Translational Science 2023;8:55–67.

50. Gavin KM, Seals DR, Silver AE, Moreau KL. Vascular endothelial estrogen receptor alpha is modulated by estrogen status and related to endothelial function and endothelial nitric oxide synthase in healthy women. J Clin Endocrinol Metab 2009;94:3513–3520.

51. Chen Z, Yuhanna IS, Galcheva-Gargova Z, Karas RH, Mendelsohn ME, Shaul PW. Estrogen receptor alpha mediates the nongenomic activation of endothelial nitric oxide synthase by estrogen. J Clin Invest 1999;103:401–406.

52. Favre J, Vessieres E, Guihot A-L, Proux C, Grimaud L, Rivron J, Garcia MCL, Réthoré L, Zahreddine R, Davezac M, Fébrissy C, Adlanmerini M, Loufrani L, Procaccio V, Foidart J-M, Flouriot G, Lenfant F, Fontaine C, Arnal J-F, Henrion D. Membrane estrogen receptor alpha (ERα) participates in flow-mediated dilation in a ligand-independent manner. eLife 2021;10:e68695.

53. Arnal JF, Lenfant F, Metivier R, Flouriot G, Henrion D, Adlanmerini M, Fontaine C, Gourdy P, Chambon P, Katzenellenbogen B, Katzenellenbogen J. Membrane and Nuclear Estrogen Receptor Alpha Actions: From Tissue Specificity to Medical Implications. Physiol Rev 2017;97:1045–1087.

54. Smirnova NF, Fontaine C, Buscato M, Lupieri A, Vinel A, Valera MC, Guillaume M, Malet N, Foidart JM, Raymond-Letron I, Lenfant F, Gourdy P, Katzenellenbogen BS, Katzenellenbogen JA, Laffargue M, Arnal JF. The Activation Function-1 of Estrogen Receptor Alpha Prevents Arterial Neointima Development Through a Direct Effect on Smooth Muscle Cells. Circ Res 2015;117:770–778.

55. Gavin KM, Seals DR, Silver AE, Moreau KL. Vascular Endothelial Estrogen Receptor α Is Modulated by Estrogen Status and Related to Endothelial Function and Endothelial Nitric Oxide Synthase in Healthy Women. The Journal of Clinical Endocrinology & Metabolism 2009;94:3513–3520.

56. Shearman AM, Cooper JA, Kotwinski PJ, Miller GJ, Humphries SE, Ardlie KG, Jordan B, Irenze K, Lunetta KL, Schuit SCE, Uitterlinden AG, Pols HAP, Demissie S, Cupples LA, Mendelsohn ME, Levy D, Housman DE. Estrogen Receptor α Gene Variation Is Associated With Risk of Myocardial Infarction in More Than Seven Thousand Men From Five Cohorts. Circulation Research 2006;98:590–592.

57. Shearman AM, Cupples LA, Demissie S, Peter I, Schmid CH, Karas RH, Mendelsohn ME, Housman DE, Levy D. Association between estrogen receptor alpha gene variation and cardiovascular disease. Jama 2003;290:2263–2270.

58. Sanchez AM, Flamini MI, Zullino S, Gopal S, Genazzani AR, Simoncini T. Estrogen receptor-{alpha} promotes endothelial cell motility through focal adhesion kinase. Mol Hum Reprod 2011;17:219–226.

59. Simoncini T, Scorticati C, Mannella P, Fadiel A, Giretti MS, Fu X-D, Baldacci C, Garibaldi S, Caruso A, Fornari L, Naftolin F, Genazzani AR. Estrogen Receptor α Interacts with Gα13 to Drive Actin Remodeling and Endothelial Cell Migration via the RhoA/Rho Kinase/Moesin Pathway. Molecular Endocrinology 2006;20:1756–1771.

60. Spyridopoulos I, Sullivan AB, Kearney M, Isner JM, Losordo DW. Estrogen-Receptor–Mediated Inhibition of Human Endothelial Cell Apoptosis. Circulation 1997;95:1505–1514.

61. Fortini F, Vieceli Dalla Sega F, Caliceti C, Aquila G, Pannella M, Pannuti A, Miele L, Ferrari R, Rizzo P. Estrogen receptor β–dependent Notch1 activation protects vascular endothelium against tumor necrosis factor α (TNFα)-induced apoptosis. Journal of Biological Chemistry 2017;292:18178–18191.

62. van Wezenbeek J, Groeneveldt JA, Llucià-Valldeperas A, van der Bruggen CE, Jansen SMA, Smits AJ, Smal R, van Leeuwen JW, Remedios Cd, Keogh A, Humbert M, Dorfmüller P, Mercier O, Guignabert C, Niessen HWM, Handoko ML, Marcus JT, Meijboom LJ, Oosterveer FPT, Westerhof BE, Heijboer AC, Bogaard HJ, Vonk Noordegraaf A, Goumans MJ, de Man FS. Interplay of sex hormones and long-term right ventricular adaptation in a Dutch PAH-cohort. The Journal of Heart and Lung Transplantation 2022;41:445–457.

63. Baxan N, Zhao L, Ashek A, Niglas M, Wang D, Khassafi F, Sabrin F, Dubois O, Chen C-N, Pullamsetti SS, Wilkins M, Zhao L. Deep phenotyping the right ventricle to establish translational MRI biomarkers for characterization of adaptive and maladaptive states in pulmonary hypertension. Scientific Reports 2024;14:29774.

64. Vélez-Rendón D, Zhang X, Gerringer J, Valdez-Jasso D. Compensated right ventricular function of the onset of pulmonary hypertension in a rat model depends on chamber remodeling and contractile augmentation. Pulm Circ 2018;8:2045894018800439.

65. Wu B, Zhang Z, Lui W, Chen X, Wang Y, Chamberlain AA, Moreno-Rodriguez Ricardo A, Markwald Roger R, O’Rourke Brian P, Sharp David J, Zheng D, Lenz J, Baldwin HS, Chang C-P, Zhou B. Endocardial Cells Form the Coronary Arteries by Angiogenesis through Myocardial-Endocardial VEGF Signaling. Cell 2012;151:1083–1096.

66. Bogaard HJ, Natarajan R, Mizuno S, Abbate A, Chang PJ, Chau VQ, Hoke NN, Kraskauskas D, Kasper M, Salloum FN, Voelkel NF. Adrenergic receptor blockade reverses right heart remodeling and dysfunction in pulmonary hypertensive rats. Am J Respir Crit Care Med 2010;182:652–660.

67. Kassa B, Kumar R, Mickael C, Sanders L, Vohwinkel C, Lee MH, Gu S, Poth JM, Stenmark KR, Zhao YY, Tuder RM, Graham BB. Endothelial cell PHD2-HIF1α-PFKFB3 contributes to right ventricle vascular adaptation in pulmonary hypertension. Am J Physiol Lung Cell Mol Physiol 2021;321:L675–l685.

68. Honigberg MC, Patel AP, Lahm T, Wood MJ, Ho JE, Kohli P, Natarajan P. Association of premature menopause with incident pulmonary hypertension: A cohort study. PLoS One 2021;16:e0247398.

