## Supplementary material for "Sexually dimorphic role of estrogen receptor α in preserving right ventricular endothelial integrity": All supplementary information

### Supplemental Materials and Methods

#### Rats and housing conditions

ERα loss-of-function (ERα$^{Mut}$) rats were generated as previously described[16]. Wild-type (WT) and ERα$^{Mut}$ male and female rats were used in this study. All procedures were approved by National Jewish Health Institutional Animal Care and Use Committee (Protocol #AS2846-09-24) and were adherent with the National Institute of Health guidelines for care and use of laboratory animals under the animal welfare assurance act. Rats were allowed *ad libitum* access to food and water and were housed in a facility with a 12-hour light/dark cycle. Animals were randomly assigned to experimental groups.

Because the phenotype associated with loss of functioning ERα is female-predominant (confirmed in this study), data from female animals and cells are shown in the main manuscript, while data from male animals and cells are shown in this Supplement.

#### E2 ELISA

5 ml of blood was collected through RV puncture from male and female WT and ERα$^{Mut}$ rats and centrifuged at 2500 g for 15 min. Serum was then collected from the top layer without disturbing the bottom layer. E2 ELISA (Cayman Chemical) was then performed according to manufacturer's instruction. 50 μl of serum was used in each well. E2 concentrations were calculated using manufacturer provided work sheet (https://www.caymanchem.com/analysisTools/elisa).

#### RVEC isolation

RVs were dissected from male and female rats, minced, and digested in 12-well plate with Collagenase Type IV (final concentration 2mg/ml), Dispase II (final concentration 1.2 U/ml), $CaCl_2$ (final concentration 0.9 mM) in HBSS (1ml/well) at 37°C for 1hr. Cell suspensions were resuspended every 15 min. After 1hr, cell suspensions were strained with 40 μm mesh strainer and quenched with 1ml of 15% horse serum in Ham's F10 w/ L-Glutamine. Cells were then pelleted and resuspended in 1ml of red blood cell lysis buffer, followed by adding 10 ml of quenching buffer. Then cells were positively selected using Pan-mouse IgG Dynabeads (Thermofisher) coated with mouse monoclonal anti-rat CD31 antibody (BD Sciences) and seeded into a gelatin-coated 6-well plate in EGM-2 MV (Lonza) media supplemented with Normocin (Invivogen).

**Primary cell culture**

Endothelial cell lineage was validated based on morphology, von Willebrand factor, vascular endothelial cadherin (VECAD) and CD31 staining, as well as Matrigel tube formation[27]. Cells were cultured at 37°C in a humidified atmosphere containing 5% $CO_2$ and were used in experiments to passage 8 in EGM-2 MV (Lonza). Cells were routinely tested for mycoplasma contamination (Lonza).

**Tissue homogenization**

Rat RV tissue was homogenized using Omni International tissue grinder (ThermoFisher) in ice-cold RIPA lysis buffer (ThermoFisher) containing proteinase inhibitor cocktail (EMD-Millipore-Sigma Aldrich) and PhosStop inhibitor cocktail (Roche). After homogenization, lysates were sonicated for ten 1-second pulses at 100% power and then centrifuged at 10,000 rpm for 10 min. The supernatant was saved and used as RV lysate.

**Angiogenesis PCR array**

RNA from WT and $ER\alpha^{Mut}$ RVECs was isolated using RNeasy Mini Kit (Qiagen). RNA concentrations were measured with Nanodrop (ThermoFisher). cDNA was reverse-transcribed using $RT^2$ First Strand Kit (Qiagen). $RT^2$ Profiler™ PCR Array (Rat Angiogenesis, GeneGlobe Id: PARN-024ZA-2) was performed according to the manufacturer's instructions. Gene expression was quantified with RT² SYBR Green ROX qPCR Mastermix (Qiagen) using Quantstudio 3. Results were analyzed with GeneGlobe web-based platform (Qiagen).

### Supplementary Results

**Suppl. Fig 1.**

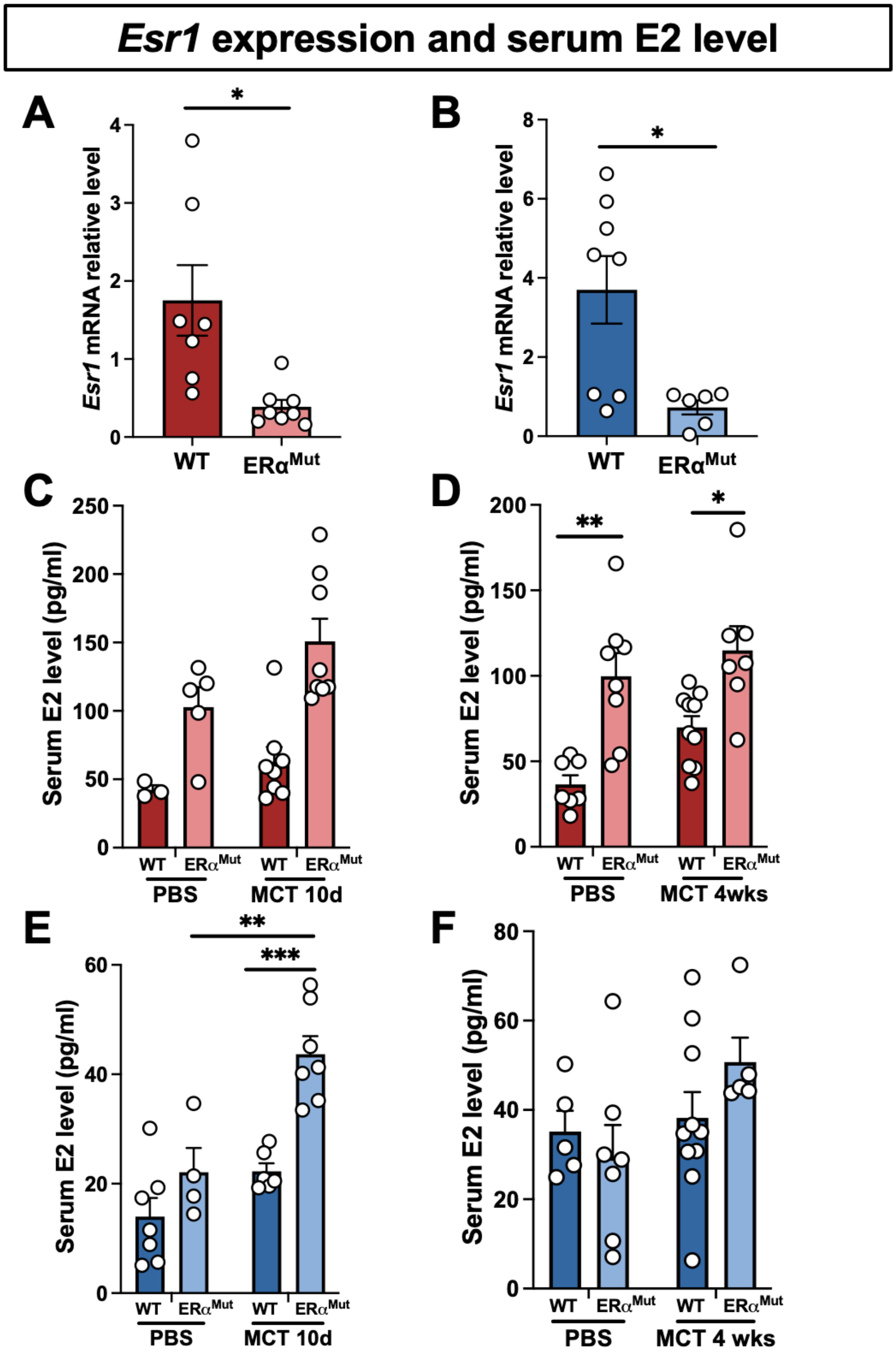

**Supplementary Figure 1. RVEC *Esr1* expression levels and serum E2 levels in female and male WT and ER$\alpha^{Mut}$ rats**. (**A, B**) Bar graphs of *Esr1* expression measured by Taqman qPCR assay in female (**A**) and male (**B**) RVECs. Relative expression level, $2^{-\Delta\Delta Ct}$ calculation. (**C, D**) Serum E2 levels in female WT and ER$\alpha^{Mut}$ rats with 10-day MCT treatment (**C**), and 4-week MCT treatment (**D**). (**E, F**) Serum E2 levels in male WT and ER$\alpha^{Mut}$ rats with 10-day MCT treatment (**E**) and 4-week MCT treatment (**F**). Each data point = one animal. * $p<0.05$, ** $p<0.01$; *** $p<0.001$ by unpaired t-test or ANOVA with Tukey's post-hoc analysis. Error bars represent mean ± SEM.

Suppl. Fig 2.

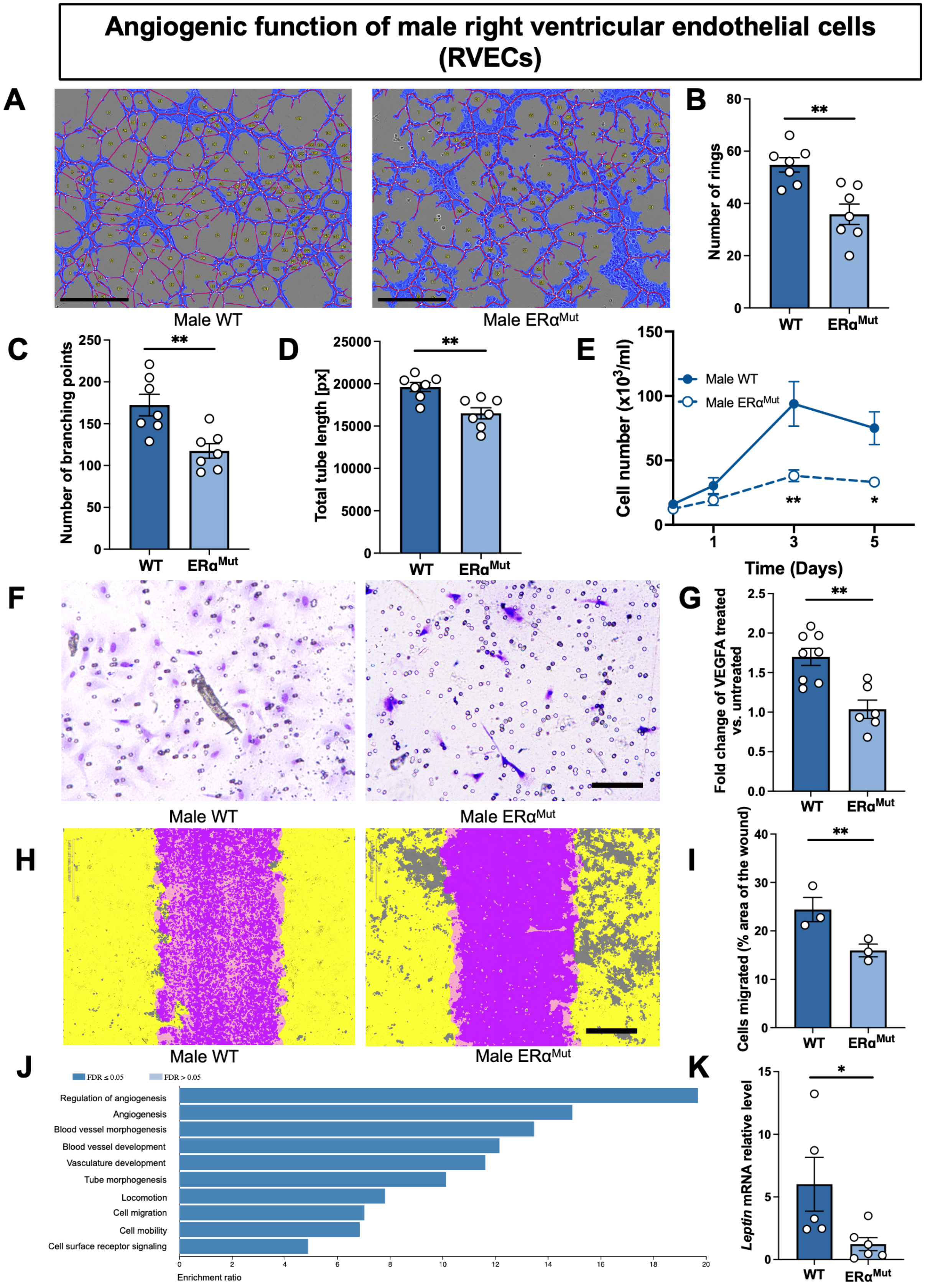


66

**Supplementary Figure 2. Loss of ERα reduces angiogenesis and migration in male RVECs.** (**A**) Representative images of ube formation assays at 10h after plating cells on Matrigel. **(B-D)** Quantifications of number of rings (**B**), numbers of branching points (**C**) and total tube length (**D**). (**E**) Growth curves of male WT and ERα$^{Mut}$ RVECs. (**F**) Transwell migration assay representative images and (**G**) quantification in male WT and ERα$^{Mut}$ RVECs at 24h. (**H**) Scratch assay representative images and (**I**) quantification in and male WT and ERα$^{Mut}$ RVECs. Purple shade represents initial wound at 24h. (**J**) GO biological process comparisons between female WT and ERα$^{Mut}$ RVECs from Qiagen rat angiogenesis microarray (n=4 animals/group, pooled). (**K**) qPCR analysis of *leptin* RNA expression of male WT and ERα$^{Mut}$ RVECs from Taqman gene expression. Each data point = one animal. * $p<0.05$, ** $p<0.01$; *** $p<0.001$ by unpaired t-test or ANOVA with Tukey's post-hoc analysis. Error bars represent mean ± SEM. Relative expression level, $2^{-\Delta\Delta Ct}$ calculation.

**Suppl. Fig 3.**

### RVEC angiogenesis PCR array

**A**

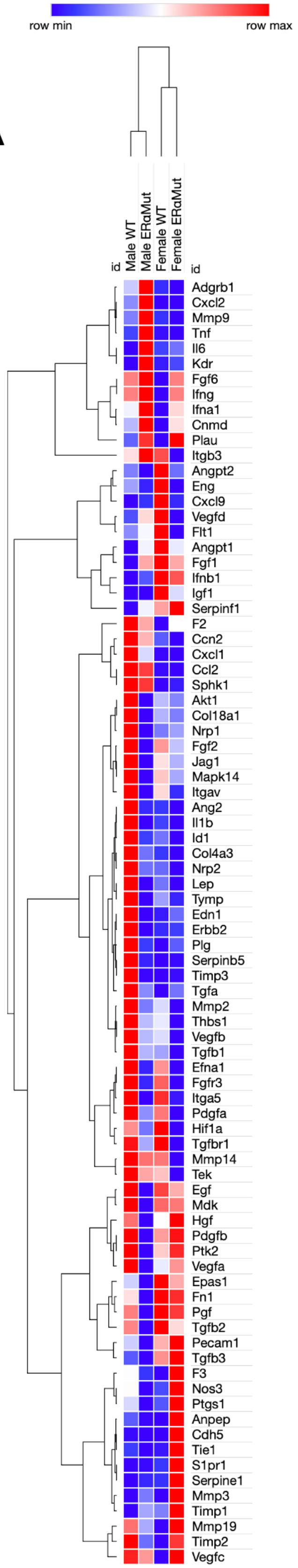


**B**

**Male WT vs ERα$^{Mut}$**

| Gene Symbol | Fold Change |
|---|---|
| Pdgfb | 12.93 |
| Lep | 11.55 |
| Il1b | 10.65 |
| Serpinb5 | 7.71 |
| Col18a1 | 4.80 |
| Il6 | -13.31 |
| Kdr | -12.11 |
| Mmp3 | -4.46 |
| Tnf | -3.45 |
| Cxcl2 | -2.82 |

**Female WT vs ERα$^{Mut}$**

| Gene Symbol | Fold Change |
|---|---|
| Cxcl9 | 4.29 |
| Lep | 3.68 |
| Il1b | 2.39 |
| Vegfd | 2.29 |
| Itgb3 | 2.22 |
| Cdh5 | -75.16 |
| Mmp3 | -14.30 |
| Tie1 | -12.75 |
| S1pr1 | -6.02 |
| Anpep | -4.12 |

**Supplementary Figure 3. Differential gene expression in male and female WT and ERα$^{Mut}$ RVECs determined by angiogenesis microarray**. (**A**) Hierarchical clustering of angiogenesis genes from male and female WT and ERα$^{Mut}$ RVECs (n=4 animals/group, pooled). (**B**) Top 5 upregulated and downregulated genes with each comparison. Hierarchical clustering is calculated by one minus Pearson correlation. Fold-Change (2^(- ΔΔCT)) is the normalized gene expression (2^(- ΔCT)) in test genes divided the normalized gene expression (2^(- ΔCT)) in housekeeping genes.

Suppl. Fig 4.

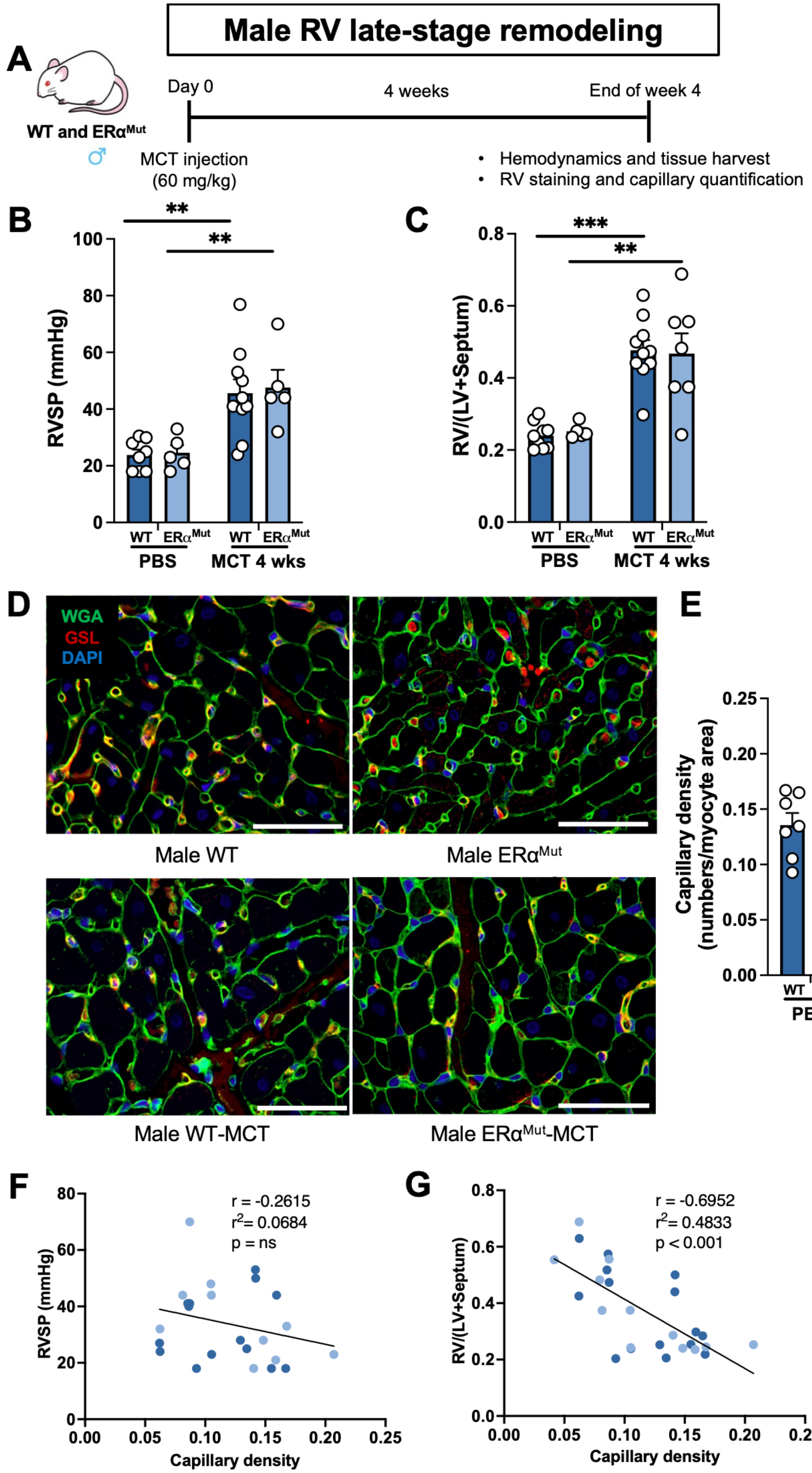


88

**Supplementary Figure 4. ERα maintains higher RV capillary density in male PH rats.** (**A**) Schematic of experimental design. (**B-E**) Effects of ERα in male rats on hemodynamics, RV hypertrophy and RV capillary density. (**B**) RVSP and (**C**) Fulton index in male WT and $ER\alpha^{Mut}$ rats treated with MCT or PBS (vehicle control). (**D, E**) Representative images of RV lectin staining and quantification as capillary per myocyte area of male WT and $ER\alpha^{Mut}$ rats. **(F, G)** Pearson correlation of capillary per myocyte area and RVSP (**F**) or Fulton Index (**G**). Dark dots indicate WT rats and light dots indicate $ER\alpha^{Mut}$ rats. WGA (Wheat Germ Agglutinin), cell membrane. GSL (Griffonia Simplicifolia Lectin), endothelial cells. Each data point = one animal. * p<0.05, ** p<0.01; *** p<0.001 by ANOVA with Tukey's post-hoc analysis. Error bars represent mean ± SEM. Scale bars, 50μm

Suppl. Fig 5.

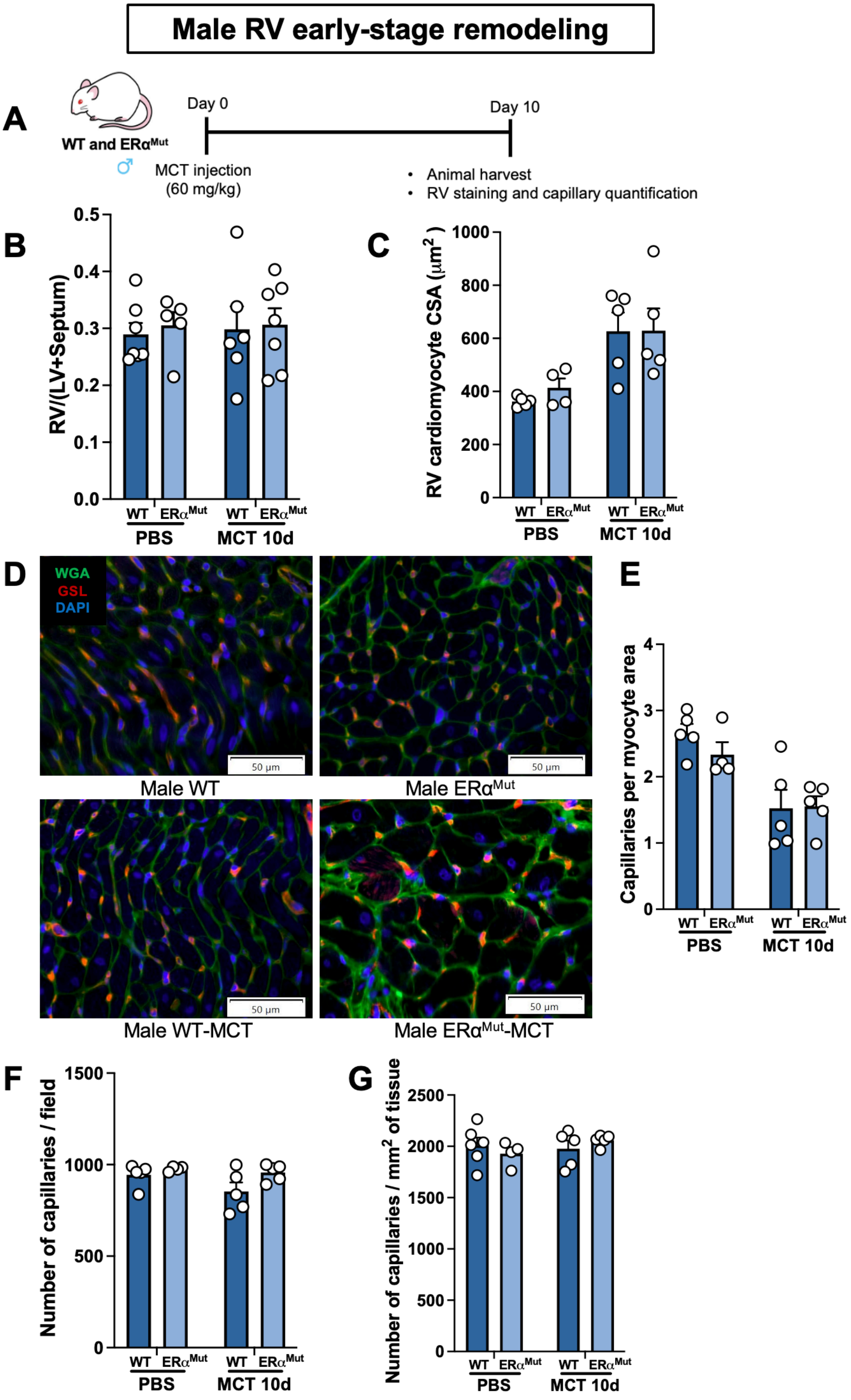


99

**Supplementary Figure 5. RV hypertrophy or RV capillary density do not differ in male WT vs ERα$^{Mut}$ rats 10 days after MCT injection.** (**A**) Schematics of experimental design. (**B-G**) Effects of ERα in male rats on RV hypertrophy and RV capillary density. Fulton index (**B**) and RV cardiomyocyte size (**C**) in male WT and ERα$^{Mut}$ rats. (**D-G**) Representative images (**D**) and quantification (**E-G**) of RV lectin staining. WGA (Wheat Germ Agglutinin), cell membrane. GSL (Griffonia Simplicifolia Lectin), endothelial cells. CSA, cross-sectional area. Each data point = one animal. * $p<0.05$, ** $p<0.01$; *** $p<0.001$ by ANOVA with Tukey's post-hoc analysis. Error bars represent mean ± SEM. Scale bars, 50μm

Suppl. Fig 6.

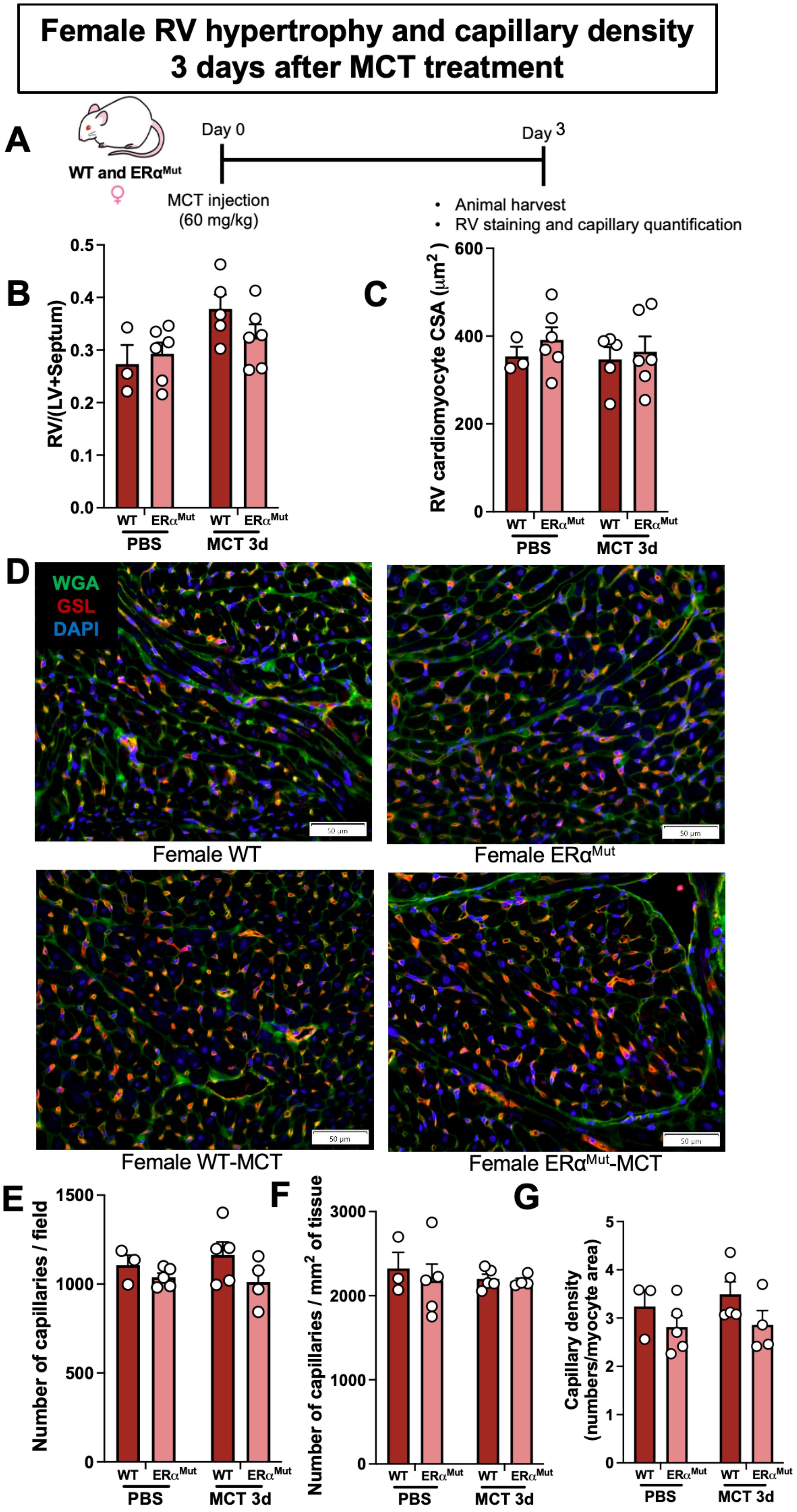


108

**Supplementary Figure 6. No capillary rarefaction is observed in female rats 3 days after MCT injection.** (**A**) Schematic of experimental design. (**B-G**) Effects of ERα in female rats on RV hypertrophy and RV capillary density. Fulton Index (**B**) and RV cardiomyocyte size (**C**) in male and female WT and $ER\alpha^{Mut}$ rats. Representative images (**D**) and quantification (**E-G**) of RV lectin staining. WGA (Wheat Germ Agglutinin), cell membrane. GSL (Griffonia Simplicifolia Lectin), endothelial cells. Each data point = one animal. * $p<0.05$, ** $p<0.01$; *** $p<0.001$ by ANOVA with Tukey's post-hoc analysis. Error bars represent mean ± SEM. Scale bars, 50μm

**Suppl. Fig 7.**

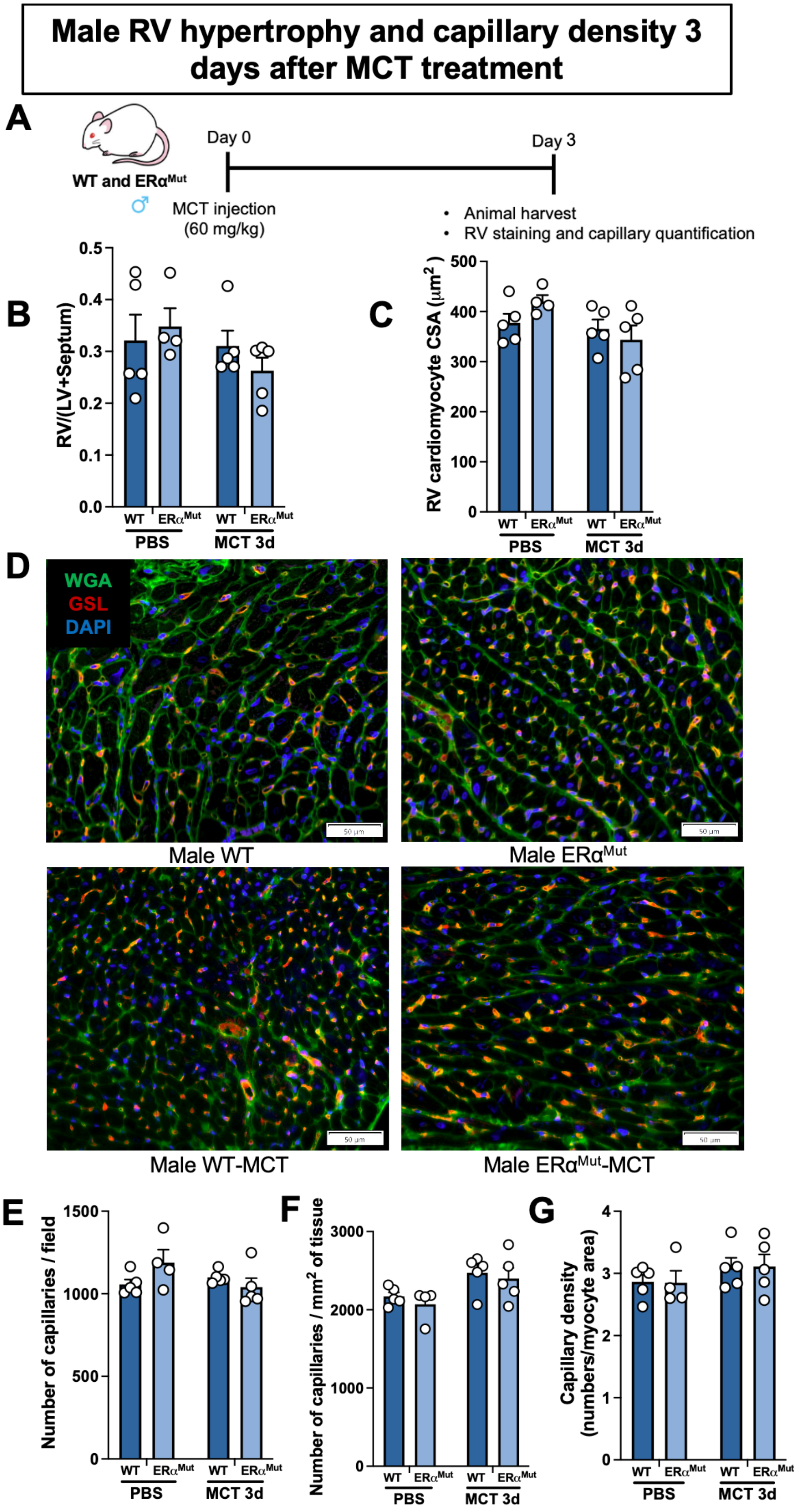


116

**Supplementary Figure 7. No capillary rarefaction is observed in male rats 3 days after MCT injection.** (**A**) Schematic of experimental design. (**B-G**) Effects of ERα in female rats on RV hypertrophy and RV capillary density. Fulton Index (**B**) and RV cardiomyocyte size (**C**) in male and female WT and $ER\alpha^{Mut}$ rats. Representative images (**D**) and quantification (**E-G**) of RV lectin staining. WGA (Wheat Germ Agglutinin), cell membrane. GSL (Griffonia Simplicifolia Lectin), endothelial cells. Each data point = one animal. * $p<0.05$, ** $p<0.01$; *** $p<0.001$ by ANOVA with Tukey's post-hoc analysis. Error bars represent mean ± SEM. Scale bars, 50μm

**Suppl. Fig 8.**

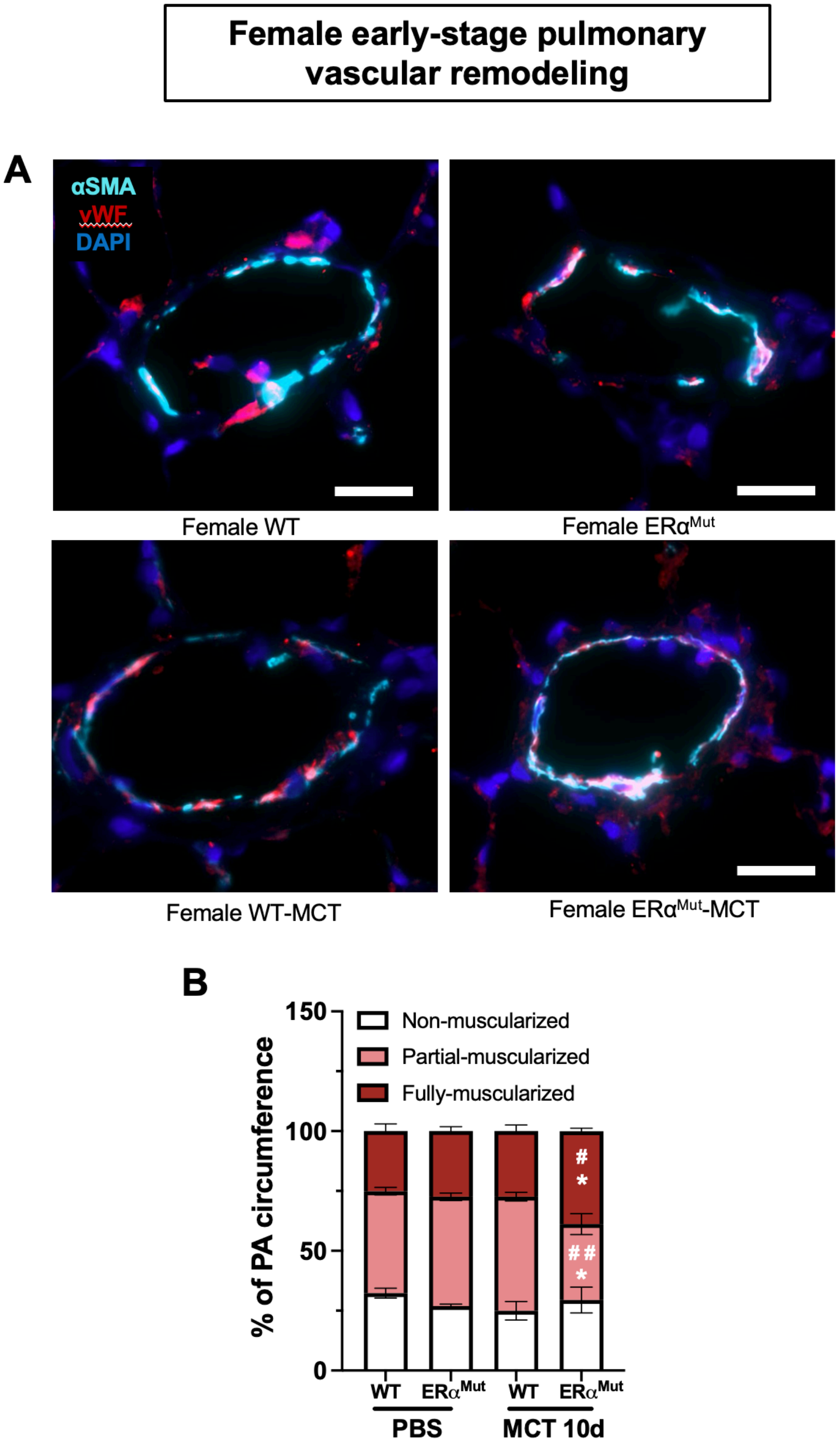
Female early-stage pulmonary vascular remodeling
A
αSMA
vWF
DAPI
Female WT
Female ERα^Mut
Female WT-MCT
Female ERα^Mut-MCT
B
150
100
50
0
% of PA circumference
Non-muscularized
Partial-muscularized
Fully-muscularized
#
*
##
*
WT
ERα^Mut
WT
ERα^Mut
PBS
MCT 10d

**Supplementary Figure 8. Loss of ERα induces pulmonary vascular remodeling in female MCT-induced PH $ER\alpha^{Mut}$ rats at day 10 after MCT administration.** Alveolar duct–associated pulmonary arteries (<200 μm) were divided into non-muscularized (α-SMA staining <25% of vessel circumference), partially muscularized (α-SMA staining 25–74% of vessel circumference), or fully muscularized (α-SMA staining ≥75% of vessel circumference) vessels. (**A**) Representative immunofluorescent images of αSMA (adventitia) and vWF (intima) staining. (**B**) Quantification of percentage of pulmonary artery muscularization. αSMA , alpha smooth muscle actin. vWF, von Willebrand factor. Each data point = one animal. ** p<0.01, compared to the same genotype with different treatment. $^{\#}$ p<0.05, compared to the different genotypes with same treatments; $^{\#\#}$ p<0.01, compared to the different genotypes with same treatments by ANOVA with Tukey's post-hoc analysis. Error bars represent mean ± SEM. Scale bars, 20μm

**Suppl. Fig 9.**

**Male early-stage pulmonary vascular remodeling**

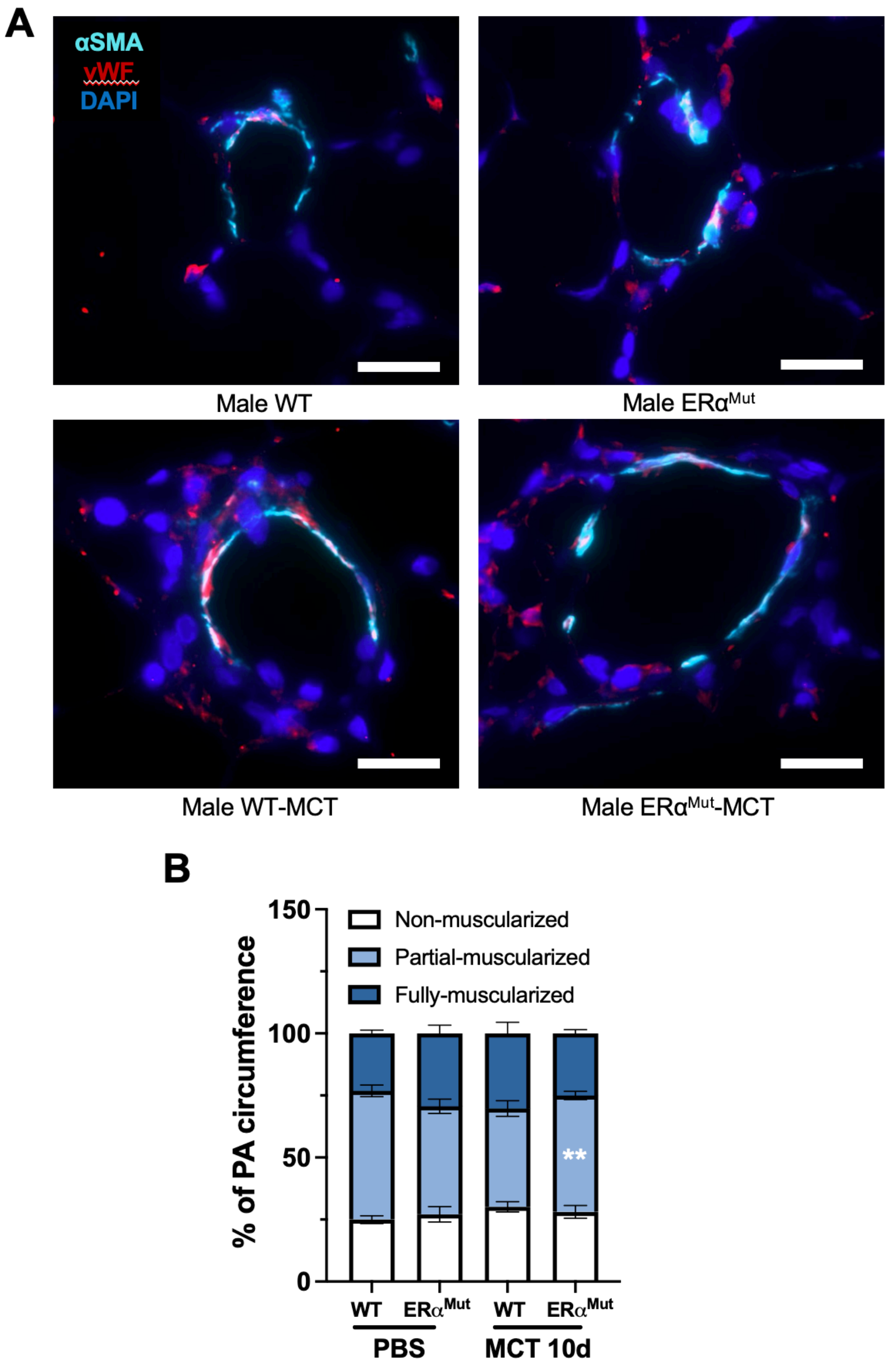

**Supplementary Figure 9. No pulmonary vascular remodeling differences were observed between male WT and ERα$^{Mut}$ rats at day 10 after MCT injection.** Alveolar duct–associated PAs (<200 µm) were divided into nonmuscularized (α-SMA staining <25% of vessel circumference), partially muscularized (α-SMA staining 25–74% of vessel circumference), or fully muscularized (α-SMA staining ≥75% of vessel circumference) vessels. (**A**) representative immunofluorescent images of αSMA (adventitia) and vWF (intima) staining. (**B**) quantification of percentage of pulmonary artery from (**A**). αSMA , alpha smooth muscle actin. vWF, von Willebrand factor. Each data point = one animal.  ** $p<0.01$, compared to the same genotype with different treatment by ANOVA with Tukey's post-hoc analysis. Error bars represent mean ± SEM. scale bars, 20µm

**Supplementary Table 1. Detailed antibody and reagent information**

| REAGENT or RESOURCE | SOURCE | IDENTIFIER |
|---|---|---|
| **Antibodies** | | |
| vWF (C-12) | Santa Cruz Biotechnology | Cat#: SC-365712 RRID:AB_10842026 |
| αSMA (D4K9N) | Cell Signaling Technology | Cat#: 19245S RRID:AB_2734735 |
| Bax (D3R2M) | Cell Signaling Technology | Cat#: 14796S RRID:AB_2716251 |
| Bcl-2 Antibody (C-2) | Santa Cruz Biotechnology | Cat#: sc-7382 RRID:AB_626736 |
| PARP | Cell Signaling Technology | Cat#: 9542S RRID:AB_2160739 |
| CD31 Mouse anti Rat, Unlabeled, Clone: TLD 3A12 | BD Biosciences | Cat#: 555025; RRID:AB395655 |
| Anti-Vinculin Antibody, clone V284 | Millipore Sigma | Cat#: 05-386 |
| ERG | Abcam | Cat#: ab92513 RRID:AB_2630401 |
| F(ab')2-Goat anti-Rabbit IgG (H+L) Cross-Adsorbed Secondary Antibody, Alexa Fluor™ 546 | Invitrogen | Cat#: A11071 |
| Goat anti-Mouse IgG (H+L) Cross-Adsorbed Secondary Antibody, Alexa Fluor™ 594 | Invitrogen | Cat#: A11005 |
| Goat anti-Mouse IgG (H+L) Highly Cross-Adsorbed Secondary Antibody, Alexa Fluor™ 488 | Invitrogen | Cat#: A11029 |
| Goat anti-Rabbit IgG (H+L) Highly Cross-Adsorbed Secondary Antibody, Alexa Fluor™ Plus 488 | Invitrogen | Cat#: A32731 |
| Donkey anti-Rabbit IgG (H+L) Highly Cross-Adsorbed Secondary Antibody, Alexa Fluor™ 647 | Invitrogen | Cat#: A31573 |
| Donkey anti-Mouse IgG (H+L) Highly Cross-Adsorbed Secondary Antibody, Alexa Fluor™ 647 | Invitrogen | Cat#: A31571 |
| Goat-anti-mouse HRP | Azure Biosystems | Cat#: AC2115 |
| Goat-anti-rabbit HRP | Azure Biosystems | Cat#: AC2114 |
| **Taqman Primers** | | |
| *Esr1* (Exon 5-6) | ThermoFisher Scientific | Rn01430446_m1 |
| *Hprt1* | ThermoFisher Scientific | Rn01527840_m1 |
| *Lep* | ThermoFisher Scientific | Rn00565158_m1 |
| **Experimental models: cell lines** | | |
| Rat: primary right ventricular endothelial cells (RVECs) | This study | N/A |
| **Experimental models: Organisms/strains** | | |

| | | |
|---|---|---|
| Rat: Sprague-Dawley | Charles River Laboratory | RRID:RGD_737891 |
| Rat: Sprague-Dawley *Esr1* $^{-/-}$ | This study | N/A |
| Rat: Sprague-Dawley *Esr1* $^{+/-}$ | This study | N/A |
| **Chemicals, peptides, and recombinant proteins** | | |
| 10x Tris/Glycine/SDS | Bio-Rad | Cat#: 1610772 |
| 10x Tris/Glycine | Bio-Rad | Cat#: 1610771 |
| Calcium chloride dihydrate | Sigma-Aldrich | Cat#: C3881 |
| BSA | Sigma-Aldrich | Cat#: A7906 |
| Crotaline | Sigma-Aldrich | Cat#: C2401 |
| Dimethyl sulfoxide | Sigma-Aldrich | Cat#: D8418 |
| Collagenase Type IV | Worthington | Cat#: LS004188 |
| Dispase II | Sigma Aldrich | Cat#: D4693-1G |
| Normocin | Invivogen | Cat#: ant-nr-1 |
| Tween 20™, Ultrapure | Thermo Scientific | Cat#: J20605.AP |
| Triton™ X-100 Surfact-Amps™ | Thermo Scientific | Cat#: 28313 |
| PhosSTOP™ | Roche | Cat#: 04906837001 |
| Protease Inhibitor Cocktail | Sigma-Aldrich | Cat#: P8340 |
| Dynal Dynabeads Pan Mouse IgG, Monoclonal | Thermo Fisher Scientific | Cat#: 11-041 |
| Recombinant Human $VEGF_{165}$ | PeproTech | Cat#: 100-20 |
| Spectra™ Multicolor Broad Range Protein Ladder | Thermo Scientific | Cat#: 26634 |
| Griffonia Simplicifolia Lectin I, Rhodamine | Vector Labs | Cat#: RL-1102-2 |
| Wheat Germ Agglutinin (WGA), Fluorescein | Vector Labs | Cat#: FL-1021 |
| DAPI Solution | Thermo Fisher Scientific | Cat#: 62248 |
| ProLong™ Gold Antifade Mountant | Invitrogen | Cat#: P36930 |
| Crystal Violet, Solution (0.1% aqueous) | Ward's Science | Cat#: 470300-938 |
| Macron™ Sodium Citrate, Dihydrate, Crystal | Macron Chemicals | Cat#: 0754-12 |
| Anhydrous Citric Acid, Granular, USP | Spectrum Chemical | Cat#: CI133 |
| **Critical commercial assays** | | |
| RNeasy Plus Mini Kit | QIAGEN | Cat#: 74134 |
| iScript cDNA Synthesis Kit | Bio-Rad Laboratories | Cat#: 1708891 |
| BCA Assay Kit | Thermo Fisher Scientific | Cat#: 23225 |
| RT² First Strand Kit | Qiagen | Cat#: 330404 |
| RT² SYBR Green ROX qPCR Mastermix | Qiagen | Cat#: 330520 |
| RT² Profiler™ PCR Array Rat Angiogenesis | Qiagen | Cat#: 330231 |
| Estradiol ELISA Kit | Cayman Chemical | Cat#: 501890 |
| MycoAlert® Mycoplasma Detection Kit | Lonza | Cat#: LT07-318 |
| GEM-X Universal 3' Gene Expression v4 4-plex | 10x Genomics | Cat#: 1000779 |
| Chromium Nuclei Isolation Kit with RNase Inhibitor | 10x Genomics | Cat#: 1000494 |
| GEM-X OCM 3' Chip Kit v4 4-plex | 10x Genomics | Cat#: 1000747 |
| **Media and buffers** | | |
| Pierce™ Blocking Buffer, Protein Free | Thermo Scientific | Cat#: 37570 |

| | | |
|---|---|---|
| Restore™ PLUS Western Blot Stripping Buffer | Thermo Scientific | Cat#: 46430 |
| RIPA Lysis and Extraction Buffer | Thermo Scientific | Cat#: 89901 |
| EBM-2 Endothelial Cell Growth Basal Medium-2 | Lonza | Cat#: CC-3156 |
| EBM™ Endothelial Cell Growth Basal Medium, Phenol Red Free | Lonza | Cat#: CC-3129 |
| EGM™ -2 MV Microvascular Endothelial Cell Growth Medium-2 BulletKit™ | Lonza | Cat#: CC-3202 |
| Corning® Fetal Bovine Serum, 500 mL, Premium, United States Origin (Charcoal Dextran Stripped) | Corning | Cat#: 35-072-CV |
| Matrigel Growth Factor Reduced (GFR) Basement Membrane Matrix, Phenol Red-free, LDEV-free | Corning | Cat#: 356231 |
| Trypsin-EDTA (0.25%), phenol red | Gibco | Cat#: 25200056 |
| HBSS | Thermo Scientific | Cat#: 88284 |
| Ham's F10 w/ L-Glutamine | Gibco | Cat#: 11-550-043 |
| Red blood cell lysis buffer | Invitrogen | Cat#: 00-4333-57 |
| EmbryoMax® 0.1% Gelatin Solution | Millipore Sigma | Cat#: ES-006-B |
| Tris Buffered Saline, 10X Solution | Thermo Fisher Scientific | Cat#: BP2471 |
| Citrate buffer (for antigen retrieval) | This study | 89ml of 0.1 M Sodium Citrate and 19ml of 0.1M citrate acid adding up to 1L with MillieQ water |
| PBS | Gibco | Cat#: 10010 |
| | Corning | Cat#: 21-040-CV |
| 4x Laemmli Sample Buffer | Bio-Rad Laboratories | Cat#: 1610747 |
| SuperSignal™ West Femto Maximum Sensitivity Substrate | Thermo Scientific | Cat#: 34096 |
| **Software and algorithms** | | |
| GraphPad Prism 9 | GraphPad Software | https://www.graphpad.com |
| Image J | NIH | https://imagej.nih.gov/ij/download.html |
| Image Lab | Bio-Rad | https://www.bio-rad.com/en-us/product/image-lab-software?ID=KRE6P5E8Z |
| LabChart | ADInstruments | https://www.adinstruments.com/support/software |
| GeneGlobe | Qiagen | https://geneglobe.qiagen.com |

| | | |
|---|---|---|
| MORPHEUS | Broad Institute | https://software.broadinstitute.org/morpheus/ |
| ShinyGO 0.80 | South Dakota State University | http://bioinformatics.sdstate.edu/go/ |
| WEB-based GEne SeT AnaLysis Toolkit (WebGestalt ) | Zhang Lab at the Baylor College of Medicine | https://www.webgestalt.org/ |
| Ingenuity Pathway Analysis | Qiagen | https://digitalinsights.qiagen.com/products-overview/discovery-insights-portfolio/analysis-and-visualization/qiagen-ipa/features-analyze-with-ipa/ |
| **Other** | | |
| 4–20% Mini-PROTEAN® TGX™ Precast Protein Gels | Bio-Rad Laboratories | Cat#: 4561093 |
| Immobilon®-P PVDF Membrane | Millipore Sigma | Cat#: IPVH00010 |
| Falcon® Permeable Support for 24-well Plate with 8.0 µm Transparent PET Membrane | Corning | Cat#: 353097 |
| Incucyte® S3 Live-Cell Analysis System | Sartorius | Cat#: 4647 |
| Incucyte® Imagelock 96-well Plate | Sartorius | Cat#: BA-04856 |
| Incucyte® Woundmaker Tool | Sartorius | Cat#: 4563 |
| Mikro-Tip® pressure catheter | Millar | Cat#: 840-8160 |
| NanoDrop Spectrophotometer | Thermo Scientific | Cat#: ND-1000 |
| QuantStudio 3 real-tme PCR system | Thermo Fisher Scientific | Cat#: A28137 |
| Omni Tissue Homogenizer | OMNI International | Cat#: TH115 |
| Bright-Line™ and Hy-Lite™ Counting Chambers | Hausser Scientific | Cat#: 3100 |
| IX83 inverted microscope | Olympus Life Sciences | |
| LSRFortessa cell analyzer | BD Biosciences | |
